# Isogenic monocytes improve the responsiveness of hiPSC cardiac spheroids to cardiac stressors

**DOI:** 10.1101/2025.10.16.682827

**Authors:** Elisabeth T. Strässler, Elise L. Kessler, Emile F. van Vliet, Nino Chirico, Ellen Na, Qingqing Cai, Alain van Mil, Gabriele G. Schiattarella, Holger Gerhardt, Nicolle Kränkel, Saskia C.A. de Jager, Joost P.G. Sluijter, Ulf Landmesser

## Abstract

**Aims:** Heart failure remains a leading cause of morbidity and mortality worldwide. Suitable in vitro models to accurately replicate the pathological environment in heart failure with reduced and preserved ejection fraction (HFrEF/HFpEF) are limited, hampering mechanistic studies and drug screening. In particular, these models rarely incorporate immune cells, which play a critical role in heart failure. To address these limitations, we developed an isogenic 3D induced pluripotent stem cell (iPSC)-derived cardiac spheroid model incorporating monocytes.

**Methods and results:** Cardiac spheroids were assembled from three healthy female iPSC lines: three-cell-type (3CT) spheroids consisting of iPSC-derived cardiomyocytes, cardiac fibroblasts, and endothelial cells, and four-cell-type (4CT) spheroids additionally containing monocytes. After six days of culture, established spheroids were treated for 24 h with different known heart failure-associated triggers (glucose & tumour necrosis factor alpha (TNFα) or ischaemia with/without reoxygenation). Differences between treated and control 3CT and 4CT spheroids were investigated at the cellular, molecular, and functional levels using confocal microscopy, RNA expression (qPCR and RNA sequencing), protein secretion using proximity extension assay technology (Olink), and functional analyses of beating rate, contraction, and relaxation.

The results confirmed successful monocyte integration in 4CT spheroids, and only spheroids with monocytes (4CTs) exhibited changes in beating rate and relaxation duration upon stimulation, highlighting the necessity of incorporating immune cells to successfully mimic heart failure-associated functional changes. Along with a more pronounced global transcriptomic treatment response and inflammatory changes, additional transcriptomic alterations previously linked to heart failure in patients, as well as changes in metabolism, ion channels, and extracellular matrix pathways, were observed in 4CT compared with 3CT spheroids.

**Conclusion:** We showed that immune cell incorporation enhances the functional and transcriptional responses of engineered cardiac tissue to relevant heart failure triggers in vitro and is essential for future studies to elucidate the cellular crosstalk and pathomechanisms.

**Translational perspective:** Heart failure continues to be a predominant cause of morbidity and mortality, necessitating the development of innovative therapeutic strategies, particularly in light of the rising prevalence of obesity and diabetes mellitus. We introduced an isogenic in vitro spheroid model comprising iPSC-derived cardiomyocytes, cardiac fibroblasts, endothelial cells, and monocytes to examine the effects of heart failure-associated triggers on cardiac tissue. Our findings indicate that spheroids incorporating monocytes exhibit a more pronounced response to heart failure-associated triggers and demonstrate greater differential transcriptional and functional responses than spheroids lacking immune cells. This model

## 1. Introduction

Heart failure (HF) is a major cause of morbidity and mortality. Both HF with reduced ejection fraction (HFrEF) and HF with preserved ejection fraction (HFpEF) impose an exponentially rising burden on healthcare systems, driven by an ageing population and the increasing prevalence of comorbidities often linked to chronic inflammation.^1^ While systolic and diastolic dysfunction have traditionally been emphasised in HFrEF and HFpEF, respectively, this distinction is reductive. Both entities share overlapping features of impaired relaxation, contractility, and reserve capacity. A unifying denominator across these syndromes is the presence of early systemic inflammation, making it an attractive research target for prevention and future mechanistic therapeutic strategies. Among inflammatory drivers, sustained activation of monocytes (MCs) and macrophages (MΦs) emerges as a key pathway fuelling vascular and cardiac injury, ultimately promoting maladaptive remodelling.^2^

To investigate the underlying mechanisms of HF, many animal models have been developed over the years but concerns about their translatability due to their non-human origin are rising, demonstrated by frequent failure in late stages of pharmacological trials.^3^ Furthermore, animal models, eg for HFpEF, are diverse and highly dependent on comorbidities and triggers, while most animal species differ in their immune system and cell types compared to humans.^4^ Nowadays, human 3D in vitro models, including spheroids, organoids, microtissues, and organs-on-a-chip^5–8^ are seen as potential alternatives to animal models as also officially stated for the first time in the FDA Modernization Act 2.0 from 2022 and Directive 2010/63/EU from the European Union from 2010 and updated in 2019.^9,10^

However, it is striking that in many multicellular models, human and animal cells are combined, or cells from multiple donors are used. The clinical relevance of the multi-donor or multi-species environment is questionable; moreover, it often lacks the incorporation of inflammatory cells, despite their demonstrated role in HF development and pathophysiology.^11^ Additionally, recently developed spheroid models have included various cell types but often still lack the inclusion of innate immune cells.^3,12,13^ In contrast, the importance of monocytes for spheroid assembly and CM function has been demonstrated.^14,15^ We hypothesised that the addition of monocytes improves the ability to mimic diastolic and systolic dysfunction, as defined by relaxation and HFpEF-related proteome and transcriptome changes. Therefore, we generated a novel isogenic multicellular 3D cardiac in vitro model. This novel model comprises four cell types (4CTs): iPSC-derived cardiomyocytes (CMs), endothelial cells (ECs), cardiac fibroblasts (FBs), and monocytes (MCs). As a control, spheroids containing only three cell types (3CTs), namely CMs, ECs, and FBs, were also generated. To investigate the effect of heart failure-associated triggers on these 4CT and 3CT spheroids, we treated the spheroids with inflammatory/metabolic stimuli (TNFα & glucose), often associated with relaxation problems in humans, or hypoxia with/without reoxygenation (0.5 % O_2_ for 6 h ± normoxic conditions for 16 h) to mimic ischaemic pathophysiology, often leading to contractile dysfunction in humans.

## 2. Methods

To generate human multicellular 3D cardiac spheroids, iPSCs were differentiated into iPSC-derived CMs, FBs, ECs, and MCs, which were then assembled into cardiac spheroids (**Figure 1**). All experiments were conducted according to the criteria of the Code of Proper Use of Human Tissue in Germany. Three healthy human quality-controlled iPSC lines were generated from peripheral blood mononuclear cells and reprogrammed as previously described (Sendai virus reprogrammed NP0141-31B, NP0143-18, and 115-4H, female).^16,17^ All donors provided informed consent under the approval of the Independent Ethics Committee of the Faculty of Medicine of the University of Cologne, Germany (Application No. DRKS00009433). The cell lines were deposited in the European Bank for induced pluripotent Stem Cells (EBiSC, https://ebisc.org/) and registered in the online registry for human iPSC lines (hPSCreg).

**Figure 1.**
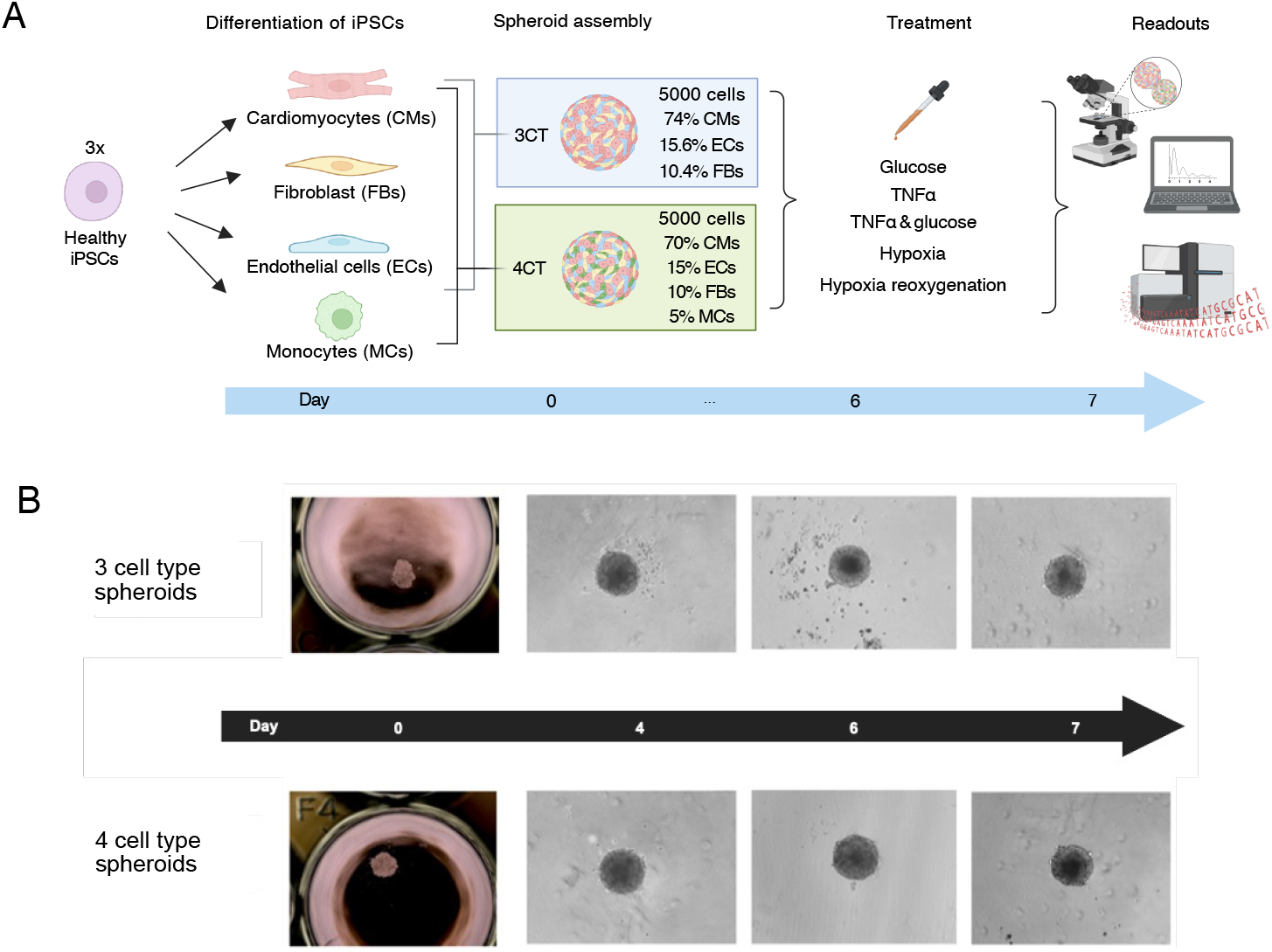
Experimental setup and assembly of three cell types (3T) and four cell types (4CT) spheroids. **A)** Four different cell types were differentiated from three healthy female induced pluripotent stem cell (iPSC) lines and assembled into cardiac spheroids consisting of three cell types (3CT) or four cell types (4CT). On day 6, spheroids were treated with either glucose, TNFα, or both TNFα and glucose to mimic left ventricular diastolic dysfunction (LVDD) and heart failure with preserved ejection fraction (HFpEF), or hypoxia and reoxygenated hypoxia to mimic heart failure with reduced ejection fraction (HFrEF) for 24 h. The next day, structural and functional alterations and differences between 3CT and 4CT were assessed. **B)** Brightfield images of 3CT (top) and 4CT (bottom) spheroids on days 0, 4, 6, and 7 show faster assembly of 4CT spheroids on day 4/6.

**Figure 2.**
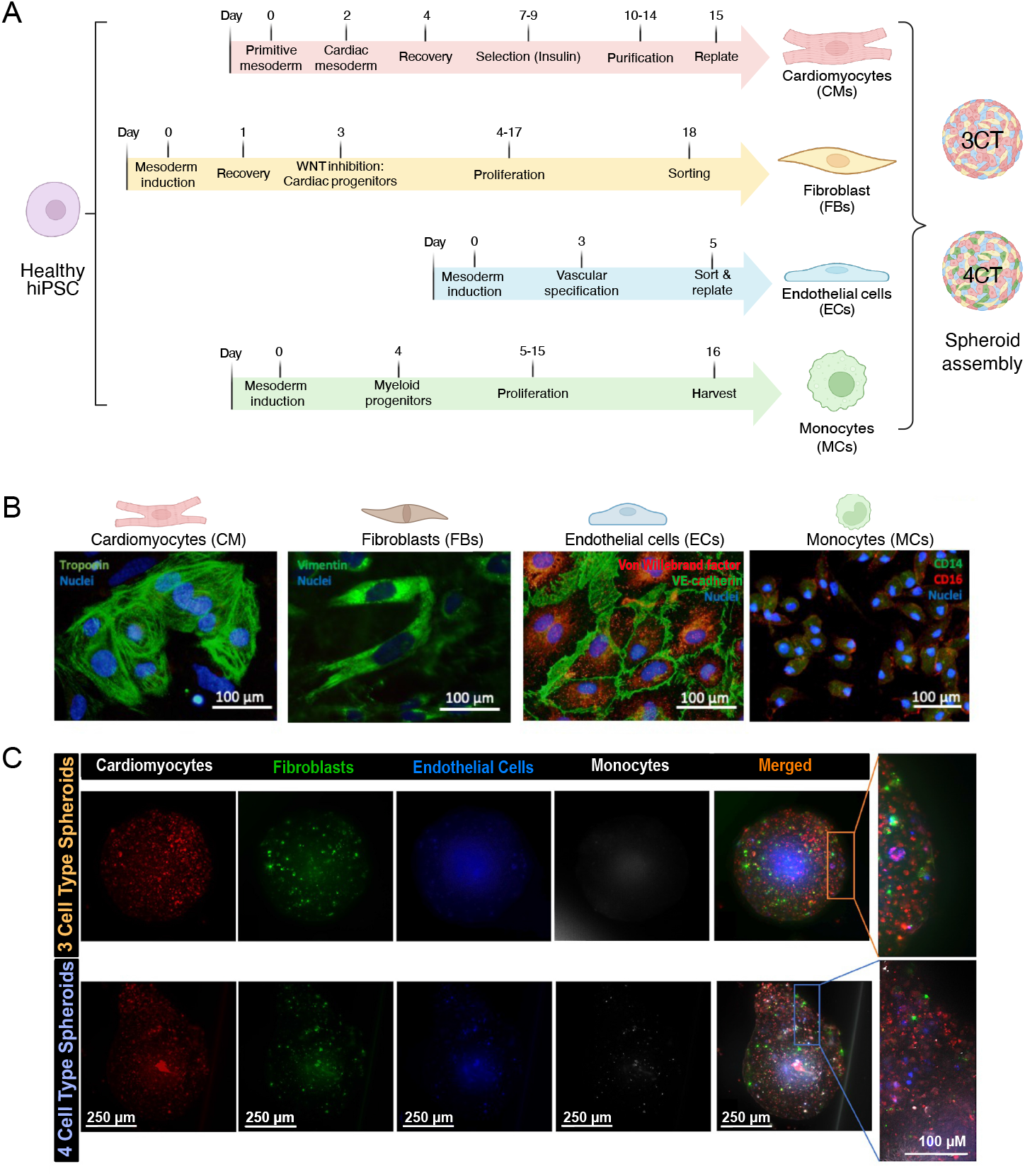
Cell type differentiation protocols, quality control, and distribution of cells in spheroids. **A)** Differentiation of iPSC-derived cardiac cardiomyocytes (CMs), fibroblasts (FBs), endothelial cells (ECs), and monocytes (MCs). Healthy iPSCs were differentiated towards the four cell types using established protocols and assembled into spheroids with or without the addition of monocytes: 4CT versus 3CT, respectively. **B)** Representative images of cell-specific fluorescent labelling of iPSC-derived cells in culture used to generate spheroids: iPSC-CMs express troponin I; iPSC-FBs express vimentin; iPSC-ECs express von Willebrand factor and VE-cadherin; and iPSC-MCs express CD14 and CD16. Blue depicts nuclei (Hoechst). Scale is 100 µm. **C)** Upper row shows a representative 3CT and lower row 4CT spheroid with pre-stained cells. iPSC-CMs are depicted in red, iPSC-FBs in green, iPSC-ECs in blue, and iPSC-MCs in white. The orange colour represents the merged picture, and the last column shows a magnification of the designated part in the square. Scale is 250 µm resp. 100 µm.

### 2.1. Differentiation of iPSC-derived cells

#### Cell culture of iPSCs

All iPSCs of all cell lines were seeded on Geltrex (Gibco A1413302)-coated well plates in Essential 8 Basal Medium (E8, Gibco A1517001) with 10 µM Rock inhibitor (Stemcell Technologies 72304). The medium was replaced daily until the start of differentiation, as shown in **Figure 1B**.

#### Differentiation of iPSC-derived cardiomyocytes

Cell culture and differentiation were performed as described previously.18 Briefly, a GiWi differentiation protocol was used.^19^ Differentiation was initiated using heparin medium (HM: DMEM/F12 (Gibco 11320033) containing 1:100 chemically defined lipid concentrate(Gibco 11905031), 213 μg/ml L-ascorbic acid (Sigma-Aldrich, A8960-5G), 1,5 IU/ml heparin (Stemcell Technologies 7980), and 1% penicillin/streptomycin (Pen/Strep, Gibco 15140148)) with 4 µM CHIR99021 (Selleck Chemicals S2924), followed by HM plus 2 µM Wnt-C59 (Tocris Bioscience 5148/10) on day 2, and then pure HM. Starting on day 7, the medium was changed every other day with insulin medium (DMEM/F12 containing 1:100 chemically defined lipid concentrate, 213 μg/ml L-ascorbic acid, 21 µg/ml human recombinant insulin (Sigma-Aldrich I9278), and 1% pen/strep). IPSC-CMs started beating around day 10, and the medium was switched to purification medium (RPMI with L-glutamine without glucose (Gibco 11879) supplemented with 3.5 μM sodium-dl-lactate (Sigma-Aldrich, L4263), 213 μg/mL L-ascorbic acid, 1% chemically defined lipid concentrate 21 µg/mL human recombinant insulin, and 1% Pen/Strep until day 15.

#### Differentiation of iPSC-derived endothelial cells

For iPSC-EC differentiation, iPSCs were seeded on Geltrex-coated 6-well plates according to the protocol described by Patsch and colleagues.^20^ After 24 h, 6 mL of N2B27 (50% DMEM/F-12 and 50% Neurobasal medium (Gibco 21103049) with 50 µM 2-mercaptoethanol (Gibco 21985023), B27 Minus Vitamin A (Gibco 12587010), N-2 Supplement (Gibco 17502001), 6 µM CHIR 99021, and 50 ng/mL bone morphogenic protein 4 (BMP4, Peprotech 120-05ET-250)) were added. On days 3 and 4, the medium was changed to 2 mL StemPro-34 medium (Gibco 10639011) with 1% Pen/Strep, 1% Glutamax (Gibco 35050061), 200 ng/µL human recombinant vascular endothelial growth factor 165 (Peprotech AF-100-20), and 2 µM forskolin (Abcam ab120058). On day 5, iPSC-ECs were purified using CD144 magnetic beads (Miltenyi Biotec 130-097-857) according to the manufacturer’s instructions, resuspended in iPSC-EC medium (Endothelium Growth Medium 2 (Promocell C-22111) with 20% FBS (Gibco 16141079), 1% Pen/Strep and 10 µM SB431542 (Selleckchem S1067)) and transferred to fibronectin (Promocell C-43060) coated T75 flasks. The iPSC-EC purity was assessed using flow cytometry (Attune NxT acoustic focusing cytometer (Life Technologies)).

#### Differentiation of iPSC-derived cardiac fibroblasts

Differentiation into iPSC-FBs was performed on Geltrex-coated plates using the protocol described by Zhang and colleagues.^21^ Medium was changed to RPMI 1640 with L-glutamine with B-27 supplement without insulin (Gibco A1895601) and 10-16 μM CHIR99021. After 24 h, the medium was changed to RPMI with B27 without insulin. On day 2, medium was changed to 2.5 mL cardiac fibroblast basal medium (high glucose DMEM (Gibco 61965026) with 500 µg/mL recombinant human albumin (Sigma A9731-1G), 0.6 µM linoleic acid (Sigma L1376-1G), 0.6 µg/mL, 50 µg/mL L-ascorbic acid (Merck A92902-25G), 3.53 µM GlutaMAX (Gibco 61965026), 1 µg/mL hydrocortisone hemisuccinate (Sigma PHR1926-500MG), 5 µg/mL rh insulin (Gibco 51500056), and 75 ng/mL basic fibroblast growth factor (bFGF, Peprotech AF-100-18B)). The medium was changed every other day until day 20, when the cells were replated in culture flasks and cultured in DMEM with 10% FBS and 1% Pen/Strep. The medium was replaced 3 times a week.

#### Differentiation of iPSC-derived monocytes

IPSC-MCs differentiation was performed as described by Buchrieser and colleagues.^22^ In short, embryoid bodies (EBs) were formed in ultra-low attachment plates (Corning 7007) in E8 medium containing 50 ng/mL BMP4, 20 ng/mL human recombinant stem cell factor (hSCF, Peprotech 300-07), 50 ng/mL VEGF, and 10 µM Rock Inhibitor (Y-27632, BD Biosciences 562822) from iPSCs to induce mesoderm. Subsequently, EBs were transferred and differentiated via the myeloid lineage from haematopoietic progenitor cells to monocytes by stimulation with 100 ng/mL human macrophage colony-stimulating factor (hM-CSF, Peprotech 300-25-100 µg) and 25 ng/mL human interleukin-3 (hIL-3; Peprotech 200-03-100) in X-Vivo-15 medium (Lonza BE02-060R) supplemented with 1% Pen/Strep, 1% GlutaMAX, and 0.1% 2-mercaptoethanol. The medium was changed every 5-7 days and monocytes were harvested.

### 2.2. Assembly and treatment of cardiac spheroids

On day 0, all cells were harvested (**Figure 1A**). iPSC-CMs, iPSC-FBs, and iPSC-ECs were first washed with PBS, detached with TryPLE Express (Gibco 12604-021), counted, centrifuged for 5 min at 300 × g, and resuspended in CO++ medium (75% low glucose DMEM (Gibco 21885025), 25% RPMI 1640 (Corning 10043CV), 10 % knockout serum replacement (KSR, Gibco 10828028), 1% Glutamax, 1 % Pen/Strep, 1 % MEM non-essential amino acids solution (Gibco 11140050), 0.1% 2-mercaptoethanol, 21 µg/ml insulin, 213 µg/ml L-ascorbic acid, 100 ng/ml VEGF 165, 50 ng/ml bFGF, and 1.5 IU/ml heparin). iPSC-MCs were harvested by collecting 2 mL of medium from three wells, counting the cells, centrifuging for 5 min at 300 × g, and resuspending in CO++ medium. Cells were combined in the following ratios for 4CT: CMs 70%, FBs 10%, ECs 15%, and MCs 5%, and for 3CT: CMs 74%, FBs 10.4%, and ECs 15.6%. In total, 5000 cells in 100 µL CO++ medium were added per well of a 96-ultra low attachment round-bottom well plate. The plate was centrifuged for 3 min at 100 × g and incubated at 37 °C with 5% CO_2_. On day 4, 50 µL of CO++ medium were replaced per well (**Figure 1B)**.

#### Treatment of cardiac spheroids

On day 6, the CO++ medium was removed, and spheroids were treated for 24 h with CO starvation medium (75% low glucose DMEM, 25% RPMI 1640, 1% KSR and 1% Pen/Strep) or CO starvation medium containing 33 mM glucose (Gibco A24940-01), 10 ng/mL TNFα (Peprotech 300-01A.100) or both TNFα & glucose mimicking metabolic pathophysiologies often leading to relaxation problems; or hypoxia (0.5 % O_2_ for 6 h) or hypoxia/reperfusion (normoxic conditions for 16 h) to mimic ischaemic pathophysiologies often leading to contraction problems.

### 2.3. Characterization of iPSC-derived cells

#### Fluorescent immunolabeling of iPSC-derived cells

For immunofluorescent labelling experiments, iPSC-CMs, FBs, ECs, and MCs were seeded on coverslips (5 × 10^4^ −2.5 × 10^6^ cells/cm^2^) and fixed using 4% paraformaldehyde (Thermo Fisher Scientific J19943.K2). Cells were permeabilised using 0.1% Triton-X-100 (Sigma-Aldrich X114-100ML) for 10 min, blocked with 10% normal goat serum (Vector Laboratories S-1000-20) for 30 min, and then incubated at 4 °C overnight with primary antibodies: CD14 (mouse anti-human, Invitrogen MA1-33348, 1:100) and CD16 (rabbit anti-human, Bios BS-6028R, 1:100) diluted in PBS. Secondary labelling was achieved using goat anti-mouse Alexa fluor-488 (Thermo Fisher Scientific A-11029, 1:400), goat anti-rabbit Alexa fluor-568 antibodies (Thermo Fisher Scientific A11036, 1:500), and 1 µg/ml Hoechst (Thermo Fisher Scientific 62249) for 4 h at room temperature.

### 2.4 Molecular, cellular, and functional readouts

#### Fluorescent labelling of the cells

To investigate the distribution of cells in the spheroids, 0.5 × 10^6^ harvested cells were resuspended in 500 µL serum-free medium (DMEM:RPMI, 3:1) with 2.5 µL staining reagent (Di-I = CM; Di-O = FB; Di-D = ECs; Di-B = MCs; (Vybrant Cell Labelling Solutions)) and incubated for 15 min at 37 °C. The cells were then washed with 5 ml of pure DMEM and centrifuged at 300 × g for 5 min. The washing step was repeated twice, and the cells were dissolved in the appropriate amount of CO++ medium to assemble spheroids consisting of 5000 cells. Spheroids were cultured as previously described. On day 7, the spheroids were fixed with 4% paraformaldehyde (Thermo Fisher J19943.K2). Images were captured using a Thunder microscope.

#### RNA sequencing of spheroids

Spheroid samples were lysed in QIAzol lysis reagent (Qiagen 79306), and total RNA was isolated using the miRNeasy micro kit (Qiagen 217084) according to the manufacturer’s instructions and sent to Single Cell Discoveries (Utrecht, The Netherlands). RNA extraction and library preparation were performed according to the CEL-seq2 protocol, with a sequencing depth of 10 million reads per sample. The R version 4.5.0. was used for the data analysis. Bulk RNA sequencing count normalisation and global differential gene expression analysis were performed using the DESeq2 package (version 1.48.1). Significantly differentially expressed genes (DEGs) between experimental groups were filtered according to their log2 fold change threshold of > ± 0.6. with adjusted p < 0.05. Heatmaps were generated using the ComplexHeatmap package (version 2.24.0). Overrepresentation analysis (ORA) was performed using the pathfindR package (version 2.4.2). Databases accessed were: KEGG - Kyoto Encyclopedia of Genes and Genomes, GO - Gene Ontology and Reactome. Plots were drawn using ggplot2 version 3.5.2, and Venn diagrams were created using the ggvenn package version 0.1.10. For comparison of the expression of maturation and immune cell markers, normalised counts were filtered for the decided markers, and expression levels were compared using the Benjamini Hochberg (BH)-adjusted Wilcoxon rank-sum test (**Figure 3C, D**).

**Figure 3.**
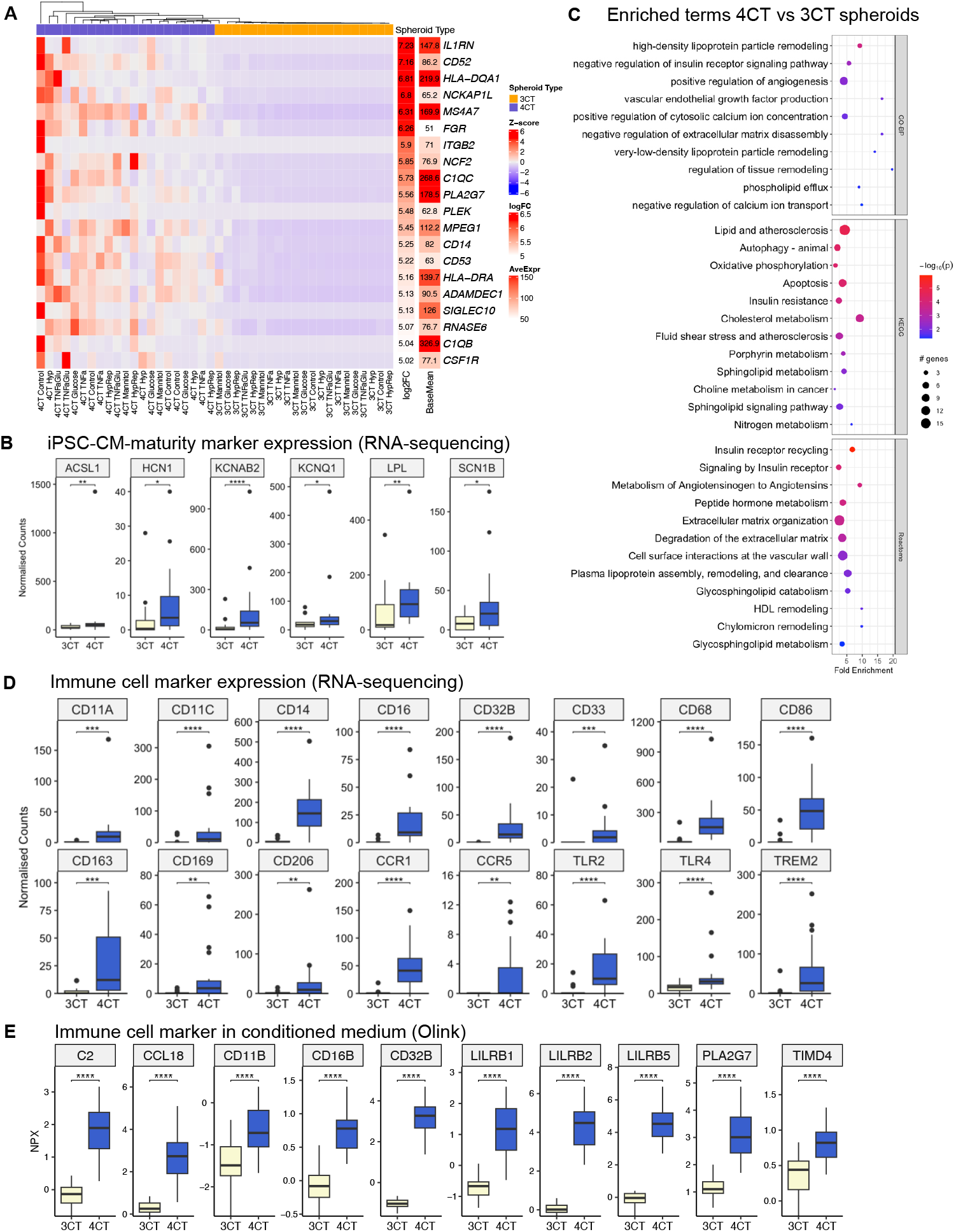
RNA-seq and targeted proteomics comparison of 4CT and 3CT spheroids. **A)** Heatmap of the top 20 differentially expressed genes (absolute log2FC > 0.6 and adjusted p < 0.05) in 4CT vs 3CT spheroids according to log_2_ fold change. **B)** Non-immunological and heart failure-associated ontology terms (GO Biological Processes, KEGG, and Reactome) for differentially expressed genes in 4CT vs 3CT. **C)** Significantly different cardiomyocyte maturity markers between 3CT and 4CT with p<0.05 in the differential gene expression analysis performed with the DESeq2 algorithm (RNA-seq normalised counts, Benjamini Hochberg (BH)-adjusted Wilcoxon rank-sum test). **D)** Gene expression of selected immune cell marker genes in 4CT vs 3CT spheroids (RNA-seq normalised counts, BH-adjusted Wilcoxon rank-sum test). **E)** Protein levels of immune cell markers in conditioned spheroid supernatant (medium after treatment) measured by Olink, presented as normalised protein expression (NPX, BH-adjusted Wilcoxon rank-sum test). N = 3 per treatment and group. Significance: * p<0.05, **p<0.01 *** p<0.001.

#### Gene expression by qPCR

Total RNA was extracted, and its quality was determined using a Nanovue (GE Healthcare). After reverse transcription of 20 ng of total RNA with the High-Capacity cDNA Reverse Transcription Kit (Applied Biosystems 4368814), Real-time PCR was performed in triplicate using the SYBR Select Master Mix (Applied Biosystems 4472918) with a ViiA 7 Real-Time PCR System (Applied Biosystems). The following primer sequences were used: GAPDH (housekeeping, forward *ggagcgagatccctccaaaat*, reverse gcaaatgagccccagccttc) CD68 (forward *caactggtgcagacagccta*, reverse *gtggtggttttgtggctctt*).

#### Secreted protein measurements by Olink

To investigate protein secretion by the spheroids, Olink cardiometabolic (Olink Proteomics AB, Uppsala, Sweden) measurements were performed in the cell-free supernatant, which was flash-frozen in liquid nitrogen on day 7, according to the manufacturer’s instructions. In short, the Proximity Extension Assay (PEA) technology utilises two antibodies conjugated to unique oligonucleotides, which facilitate hybridisation when both antibodies bind to the target protein simultaneously, bringing the oligonucleotides into proximity. The oligonucleotide sequence is then measured using qPCR. The final assay readout is presented in Normalized Protein eXpression (NPX) values, which are not absolute quantifications but an indication of the relative concentration of each analyte.

#### Beating rate and contraction and relaxation measurements

Videos and photos were taken before and after treatment on days 6 and 7 using a Moticam 5 camera in conjunction with a Motic AE2000 microscope. Beating rate and contraction measurements were performed using the Myocyter plugin for ImageJ. Contraction/relaxation measures are the relaxation duration ratio, which is the duration of the relaxation after treatment divided by the duration before treatment, and the contraction duration ratio, which is the duration of the contraction after treatment divided by the duration of contraction before treatment. For both relaxation and contraction duration ratios, the normalised values were also investigated (ratio multiplied by the beating rate) because the duration of cardiac relaxation and contraction is dependent on tissue resp. cardiac beating frequency.

#### Statistics

All data were analysed using R version 4.5.0. Statistical significance was set at p < 0.05. Before comparing two groups, the data were tested for normal distribution using the Shapiro-Wilk test. If p < 0.05, a Wilcoxon rank-sum test, also known as the Mann-Whitney U test, was performed; otherwise, a Student’s t-test was performed. P-values were adjusted for multiple testing using the Benjamini-Hochberg (BH) method, unless otherwise indicated. Olink data were analysed using the BH-adjusted Wilcoxon rank-sum test. Differential expression analysis of RNA sequencing data was performed using the DESeq2 package (version 1.48.1), as described above.

## 3. Results

### 3.1 iPSC-derived cells resemble their primary human counterparts

Cardiac spheroids can be used for a variety of studies, including testing the influence of various factors or drugs or the effect of specific gene defects (eg channelopathies). To generate 3CT and 4CT cardiac spheroids, iPSCs were differentiated into CMs, ECs, FBs, and MCs (**Figure 2A**). iPSC-CMs expressed troponin I (green), iPSC-ECs displayed von Willebrand factor (red) and VE-cadherin (green), iPSC-FBs expressed vimentin (green), and iPSC-MCs expressed CD14 (green) and CD16 (red) (**Figure 2B**).

#### Spatial distribution of iPSC-derived cells in cardiac spheroids

All spheroids consisted of 5000 cells, as this cell number showed the best density, survival, and stability (data not shown). Spheroids were seeded as follows: the 3CT spheroids were composed of 74% iPSC-CMs, 10.4% iPSC-FBs, and 15.6% iPSC-ECs. The 4CT spheroids were composed of 70% iPSC-CMs, 10% iPSC-FBs, 15% iPSC-ECs, and 5% iPSC-MCs. To investigate the spatial distribution of the cells and to demonstrate that monocytes were homogenously embedded in the 4CT spheroids, all cells were fluorescently labelled before the generation of spheroids, and images were captured after 7 days (**Figure 2C**). Here, iPSC-CMs are depicted in red, iPSC-FBs in green, iPSC-ECs in blue, and iPSC-MCs in white. We showed that cells were homogeneously (stochastically) distributed in both spheroid types and that 3CT (upper row) spheroids did not contain MCs.

Moreover, the presence of iPSC-monocyte-derived macrophages in the 4CT cardiac spheroids was confirmed using qPCR, where pooled 4CT spheroids had significantly higher CD68 expression than 3CT cardiac spheroids (*p* = 0.01, data not shown).

### 3.2 Baseline morphological, transcriptomic, and proteomics characterization of 3CT and 4CT spheroids

To characterise our spheroid model, we assessed the baseline morphology, gene expression, and secreted protein expression of 3CT and 4CT cardiac spheroids. 4CT spheroids were significantly rounder and more circular than 3CT spheroids (p<0.001; **Supplementary Figure S1A/B**). Transcriptome profiles of the spheroids were determined using RNA sequencing, and proteomics was performed on the conditioned medium using Olink targeted proteomics (96 cardiometabolic panel) (**Figure 3**).

As expected, the addition of monocytes to 4CT spheroids led to significant changes in inflammatory genes and pathways. Interestingly, we also observed differentially expressed non-inflammatory genes and pathways, suggesting that monocytes communicate with and influence other cell types in 4CT spheroids. In detail, 4CT cardiac spheroids expressed 517 significantly different genes (absolute log2FC > 0.6 and adjusted p-value < 0.05) compared to 3CTs, with **Figure 3A** depicting the heatmap of the top 20 genes.

#### Differences in cardiomyocyte maturation between 4CT and 3CT spheroids

One aspect of in vitro models containing iPSC-CMs that still needs attention is the generation of a more mature phenotype to better recapitulate adult human cardiomyocytes.^23,24^ Cardiomyocytes in the 4CT spheroids demonstrated improved maturation, as shown via enhanced expression of maturation Acyl-CoA Synthetase Long Chain Family Member 1 (ACSL1), Hyperpolarisation Activated Cyclic Nucleotide Gated Potassium Channel 1 (HCN1), potassium voltage-gated channel subfamily A regulatory beta subunit 2 (KCNAB2), Potassium Voltage-Gated Channel Subfamily Q Member 1 (KCNQ1), lipoprotein lipase (LPL), and the beta-1 subunit of voltage-gated sodium channel (SCN1B) (BH-adjusted Wilcoxon rank-sum test, p<0.05, **Figure 3C**).

#### Inflammatory and non-inflammatory transcriptomic differences between 4CT and 3CT spheroids

As expected, transcriptomics analysis showed that various immune cell-related genes and pathways were significantly more prevalent in the 4CT than in the 3CT spheroids (RNA-seq normalised counts, BH-adjusted Wilcoxon rank-sum test, **Figure 3D**). Focussing on monocyte and macrophage markers, CD14, CD16, CD11a, CD33, CD32b, CCR1, and CCR5 are characteristic monocyte markers (though some overlap with macrophages exists), and CD68, CD86, CD163, CD169, CD206, and TREM2 are more specific for tissue and other macrophage phenotypes. Shared markers include CD11c, CD32b, CCR1, CCR5, TLR2, and TLR4.^25^ Overrepresentation analysis of the differentially expressed genes between 4CT and 3CT spheroids revealed numerous non-immune-related terms relevant to heart failure2). The significantly (p < 0.05) overrepresented terms related to heart failure and cardiovascular disease in general are shown. The principal component analysis, clustered heatmap of all expressed genes, plots of cell type marker expression, and the number of differentially expressed genes for all treatments are shown in **Supplementary Figure S2**.

#### Inflammatory and non-inflammatory secreted proteomics differences between 4CT and 3CT spheroids

Analysis of the secretome of cardiac spheroids can provide critical insights into paracrine signalling and molecular (patho)physiological mechanisms, facilitating the identification of novel biomarkers and therapeutic targets. Secreted proteins were measured in cell-free supernatants using the Olink targeted proteomics cardiometabolic panel, and the most significant differences are depicted in the bar graphs (**Figure 3E**). The most differentially secreted proteins by the 4CT vs 3CT spheroids were complement component 2 (C2), C–C Motif Chemokine Ligand 18 (CCL18), Cluster of Differentiation 11b (CD11B), CD16B, CD32B, Leukocyte Immunoglobulin-Like Receptor Subfamily B Member 1 (LILRB1), LILRB2, LILRB5, Phospholipase A2 Group VII (also known as Lipoprotein-Associated Phospholipase A2, Lp-PLA2) PLA2G7, and T-Cell Immunoglobulin and Mucin Domain Containing 4, Isoform D (TIMD4). From the full list, it can be appreciated that not only inflammatory proteins were significantly different between 4CT and 3CT spheroids, but also lipoprotein catabolic, cholesterol, extracellular matrix, and endothelial cell migration processes.

### 3.3 Functional changes between 3CT and 4CT spheroids before and after heart failure associated treatment

To evaluate the 3CT and 4CT spheroids functionally, the beating rate and contraction/relaxation parameters were evaluated (**Figure 4**). At baseline, 4CT control spheroids and 3CT spheroids did not differ in beating rate (p=0.28) or relaxation duration ratio, which is comparable to patient indices of diastolic function, such as isovolumic relaxation time (IVRT), deceleration time (DT), or tau (ventricular relaxation constant) (p=0.67). They also did not differ in the contraction duration ratio, which is comparable to Left Ventricular Ejection Time (LVET) or systolic duration (p=0.88). Interestingly, the beating rate significantly increased after combined treatment with TNFα & glucose and significantly decreased after hypoxia and reoxygenated hypoxia in 4CT spheroids (**Figure 4A**). The relaxation duration ratio (after/before treatment) was significantly decreased upon treatment with TNFα & glucose and hypoxia, and significantly increased after reoxygenated hypoxia compared to hypoxia, only in 4CT spheroids (**Figure 4B**). When the relaxation duration ratio was normalised for the beating rate to distinguish whether changes in relaxation were due to intrinsic tissue effects (on relaxation itself), the hypoxia treatment remained significant compared to controls, with significant differences between reoxygenated hypoxia and hypoxia (**Figure 4C**). The contraction duration ratio results were not significantly different and are shown in **Supplementary Figure S3**.

**Figure 4.**
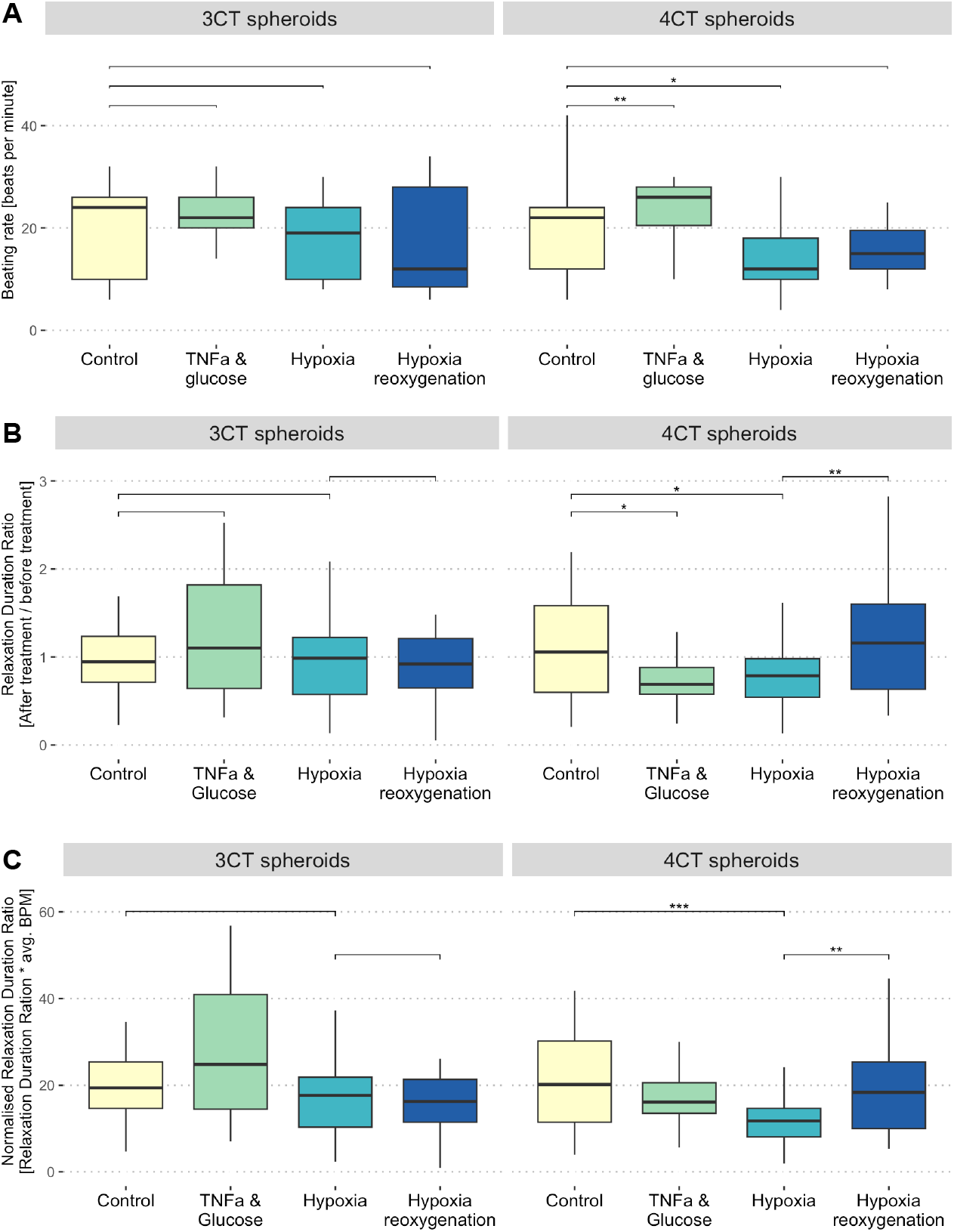
Beating rate and relaxation duration of 3CT and 4CT spheroids after several types of treatment. **A)** Beating rate (BPM) in control spheroids and after TNFα & glucose, hypoxia, and reoxygenated hypoxia treatment. **B)** Relaxation duration ratio (duration of relaxation after treatment/before treatment) in control spheroids and after TNFα & glucose, hypoxia, and reoxygenated hypoxia treatment. Relaxation duration ratio (RDR) < 1 indicates faster relaxation after treatment, whereas RDR > 1 indicates slower relaxation. The relaxation duration ratio is comparable to patient indices of diastolic function, such as isovolumic relaxation time (IVRT), deceleration time (DT), or tau (ventricular relaxation constant). **C)** Normalised diastolic duration (RDR * average BPM) to distinguish whether changes in relaxation are due to intrinsic tissue effects (on relaxation itself) as opposed to secondary effects from changes in beating frequency. Statistics: Kruskal-Wallis, followed by t-test or Wilcoxon rank-sum test, depending on p<0.05 in the Shapiro-Wilks test. ns not significant, *p<0.05, **p<0.01, ***p<0.001. N = 3 per treatment per group.

### 3.4 Transcriptomic and proteomics characterization treated 4CT spheroids

Our data indicate that the incorporation of monocytes into cardiac spheroids (4CT) leads to more pronounced functional changes than spheroids lacking monocytes (3CT), illustrating the critical role of inflammatory cells in HF-associated responses. Next, we investigated the transcriptomic differences upon TNFα & glucose stimulation or hypoxia and reoxygenation exposure in spheroids (**Figure 5**). Compared to the control, 299 genes were differentially expressed after TNFα & glucose treatment **(Figure 5A)**, 50 genes were differentially expressed after hypoxia (**Figure 5B)**, and 174 genes were differentially expressed after subsequent reoxygenation (**Figure 5C**). The small overlap between the DEGs of the 3CT and 4CT spheroids emphasises the importance of immune cell presence in the transcriptomic treatment response. A detailed description of the genes is shown in the volcano plots in **Figure 5G, I, and K**, respectively (sig. cutoff abs(log2FC) > 0.6 & p.adj < 0.05). To investigate the related pathways, an overrepresentation analysis was performed. Reactome terms are shown for the DEGs; hits included cellular senescence, as well as FGFR1, TNF, EGFR, and SMAD signalling (**Figure 5H**). After hypoxia, pathways related to IFNG, CD209, and RAF/MAP signalling were activated (**Figure 5J**). After reoxygenated hypoxia, pathways associated with NRAGE, death receptor, and Hippo signalling were activated (**Figure 5L**). When focusing on HFpEF-related transcriptomic changes (**Figure 5D, E, and F**), 4CT spheroids treated with TNFα & glucose showed almost quadruple the number of DEGs compared to 3CT spheroids (DEG = 37 vs 7, respectively). Next, 4CT spheroids that underwent reoxygenation showed larger differences in the number of differentially expressed genes than 3CT spheroids without monocytes (DEG = 32 versus 1, respectively). Overall, 4CT spheroids better recapitulated clinically observed HFpEF-associated gene expression changes, as demonstrated by the higher number of matched differentially expressed HFpEF-associated genes.

**Figure 5.**
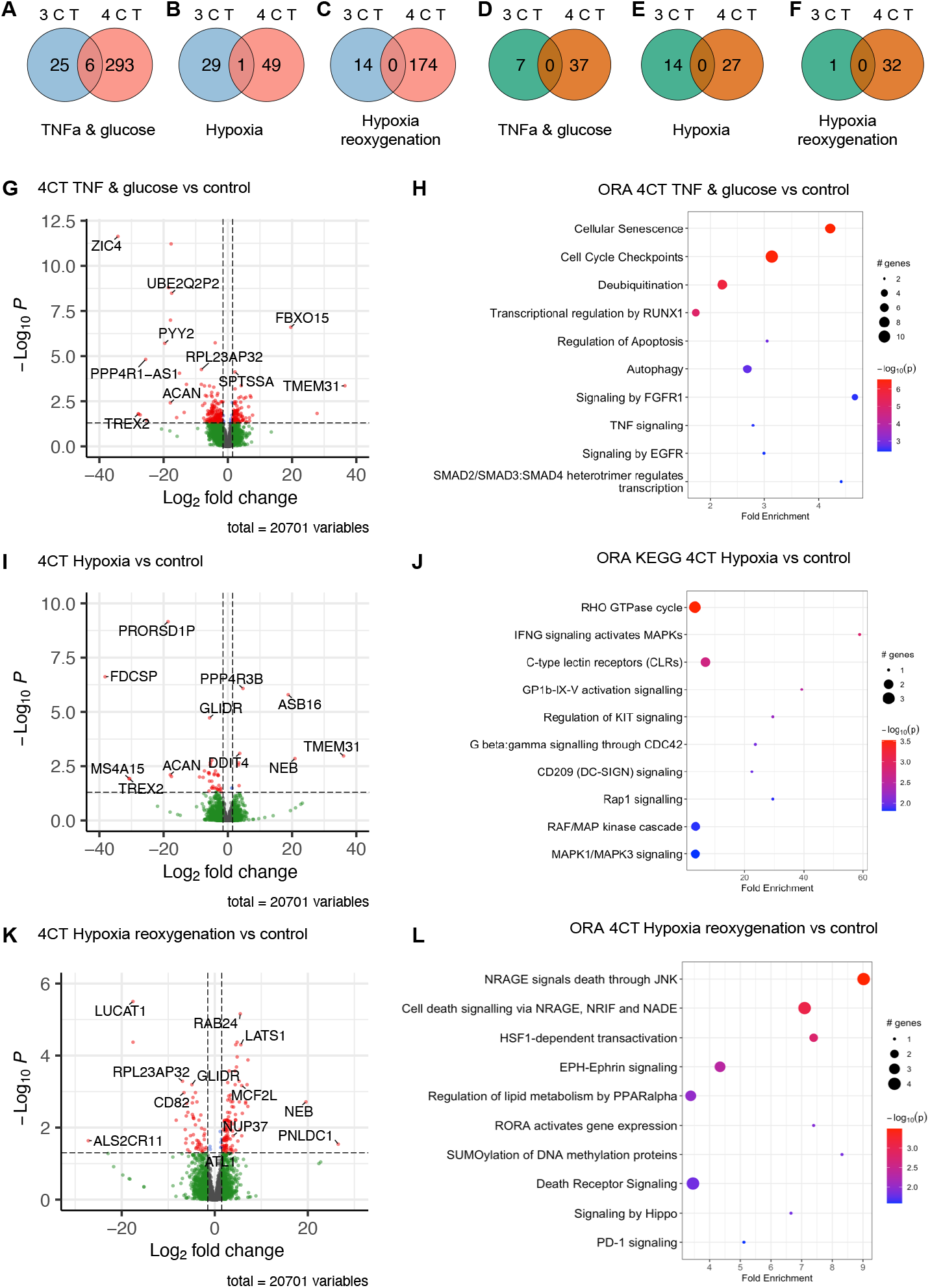
Effects of treatment on the spheroid transcriptome. *Top left:* Venn diagrams showing the number of differentially expressed genes between 3CT and 4CT spheroids under different treatment conditions. **A**) TNFα [10 ng/ml] & glucose [33mM] treatment vs control, **B)** hypoxia (0.5% O_2_ for 6 h) vs control treatment, and **C)** reoxygenated hypoxia (6 h 0.5% O_2_ followed by 16 h normoxia) vs control. *Top right:* Venn diagrams showing the number of differentially expressed HFpEF genes between 3CT and 4CT spheroids under different treatment conditions. In total, a list of 648 HFpEF-related genes was created based on the literature (Das 2019). **D**) TNFα [10 ng/ml] & glucose [33mM] treatment vs control, **E)** Hypoxia (0.5% O_2_ for 6 h) vs control treatment, and **F)** reoxygenated hypoxia (6 h 0.5% O_2_ followed by 16 h normoxia) vs control. **G)** Vulcano plot of transcriptomic changes in 4CT spheroids showing differentially expressed genes (DEGs) after TNFα & glucose treatment vs control, **H)** together with their associated Reactome terms (GO-BP). **I)** Volcano plot of DEGs in 4CT spheroids after hypoxia (0.5% O_2_) vs control, **J)** together with their associated Reactome. **K)** Volcano plot of DEGs in 4CT spheroids after reoxygenated hypoxia treatment vs control, **L)** together with their associated Reactome terms. Differential gene expression analysis was performed with DESeq2 using RStudio and Bioconductor with a cutoff of abs(log2FC) > 0.6 and an adjusted p-value < 0.05. N = 3 per treatment and group.

## 4. Discussion

This study introduces a novel isogenic 3D cardiac spheroid model to investigate the response to heart failure-associated stimuli (TNFα & glucose, hypoxia, and reoxygenated hypoxia) in the absence or presence of monocytes/macrophages. Our 3D spheroids containing monocytes (4CT) demonstrate a novel approach to modelling traits associated with heart failure in vitro by including the interaction with monocytes.

The incorporation of monocytes into the 4CT spheroids created rounder and more circular spheroids and significantly increased the maturation of iPSC-CMs **(Supplementary Figure S1 and Figure 3B)**. More mature iPSC-CMs more closely mimic the situation in human heart tissue and improve translatability.^26–29^ The monocytes in the 4CT spheroids significantly improved the model’s response to heart failure-associated stimuli. The 4CT spheroids exhibited more transcriptomic and proteomic changes than the 3CT spheroids. In addition to the expected pathways associated with inflammation (**Figure 3A, D, and E**), pathways associated with metabolism (eg cholesterol metabolism, insulin signalling, and oxidative phosphorylation), electrophysiology (eg calcium ion transport), and extracellular matrix organisation were overrepresented (**Figure 3B**). In addition, the expression of HFpEF-associated genes was increased in 4CTs compared to that in 3CT spheroids without monocytes under all treatment conditions (**Figure 5D, E, and F**). No overlap was observed in HFpEF-associated genes between 3CT and 4CT, indicating the activation of distinct cellular and molecular programs in the two spheroid types. The presence of increased gene expression is not only related to inflammation but also to other disease drivers, underscoring the critical role of immune cells in modelling cardiovascular (patho-)physiology. Moreover, 4CT spheroids exhibited functional characteristics similar to those measured in vivo, such as variations in beating rate (corresponding to heart rate) and relaxation duration, roughly corresponding to patient indices of diastolic function, such as isovolumic relaxation time (IVRT), deceleration time (DT), or tau (ventricular relaxation constant) (**Figure 4**), which 3CT spheroids did not demonstrate.

The non-immune-related changes in 4CT spheroids highlight similarities and translatability to heart failure, especially to HFpEF, where preexisting metabolic conditions (eg diabetes) and fibrotic extracellular matrix changes leading to increased cardiac stiffness are the primary drivers of disease progression.^30^ Some functional changes due to HFpEF-mimicking treatments were also shown in CM-only spheroids recently^14,31–33^ but the authors themselves highlighted the lack of non-CMs as a limitation. Our study also provides insights into modelling diastolic dysfunction, as relaxation time decreases acutely after hypoxia in this condition.^34^ An increased heart rate is associated with increased stress and metabolic disease in patients,^35^ whereas an increased relaxation time is one of the hallmarks of diastolic dysfunction.^36–38^ Therefore, we propose that this model can be applied as an early in vitro tool in heart failure research, especially for personalised medicine strategies and pharmacological evaluation.

The FDA Modernization 2.0 Act of 2022 concluded that human iPSC-derived models are promising for improving pre-clinical drug development and clinical trial pipelines.^9^ Our observations suggest that the inclusion of immune cells, such as MCs and MΦs, enables more accurate modelling of alterations observed in human cardiac tissue associated with heart failure, leading to increased translatability and efficiency of future pre-clinical approaches. The role of the innate immune system has been a topic of interest during the last decade with more clinical data supporting its relevance in heart failure; aiding phenotyping and phenogrouping of patients with HFpEF based on inflammatory risk.^39,40^ Inflammation has also emerged as a desirable drug target for CVDs in general^41^ and HFpEF in particular^42,43^ and (residual) inflammation is seen as one of the greatest risk factors for mortality.^44,45^

Despite the general acknowledgement of the role of inflammation in the pathophysiology of HF, inflammatory cells are not a default component of ex vivo models.^12^ The implementation of inflammatory cells and the resulting improved translatability of our model are clear strengths of our new 3D model. As is the improved maturation of iPSC-CMs, which is a major challenge in current cardiac in vitro research.^29,46^ Our findings of improved iPSC-CM maturation in 4CT spheroids align with other studies finding comparable results.^15,26,47^ We also found *KCNQ1* to be significantly increased in 4CTs, a potassium channel responsible for the termination of the action potential and normally not measurable in immature iPSC-CM.^48^

In vitro models for investigating the underlying cardiac pathophysiologies are on the rise to decrease the use of laboratory animals and better mimic human diseases. However, many 3D cardiac in vitro models use primary cells from various donors or even species, limiting their transferability to patients or personalised medicine.

Finally, our model is especially relevant in light of the increasing rates of heart failure in Western countries, with HFpEF now being more common than HFrEF.^1^ The Western diet, which is glucose-rich and often associated with chronic inflammation and diabetes (such as mimicked with our TNFα & glucose treatment), has been shown to cause HFpEF^49^, which is also supported by the great therapeutic effect of sodium-glucose cotransporter 2 (SGLT2) inhibitors in the clinical setting.^50^ Investigating various triggers and risk factors for HFpEF in vitro in combination with the relevant cell types will hopefully improve preclinical HFpEF research dramatically in the future, and our data will enable more researchers to learn from it to accelerate the preclinical work and finding of new treatment options.

## Limitations and future perspectives

While our data have provided valuable insights into the molecular mechanisms underlying heart failure and offered a new 3D cardiac in vitro model to investigate this, there are several limitations to this study that should be considered. One limitation is the small number of spheroids available for bulk RNA-seq, which may not provide a complete picture of the cellular response to stimuli and does not offer single-cell resolution. For instance, while no changes in fibrosis-associated gene expression were observed in this study, it is possible that other mechanisms may be involved in fibrosis development, or that the timing was optimal to detect changes in RNA expression. This could be mitigated by introducing other stimuli as treatment, such as pro-fibrotic cytokines (eg transforming growth factor beta (TGF-β) or IL-6) or neurohumoral and metabolic stressors (eg high glucose or norepinephrine). Future studies should aim to increase the sample size for RNA-seq and switch to single-cell RNA-seq to improve the statistical power and robustness and gain cell type-specific insights. Another limitation is that only a specific set of stimuli was tested in this study. Other stimuli, as well as established HF treatments (eg SGLT2 inhibitors such as dapagliflozin, interleukin-1 receptor antagonists such as anakinra, or beta-blockers), should be tested to better understand the cellular response to different triggers. Additionally, incorporating electrophysiology into the readouts could provide further understanding of the physiological responses of cells to stimuli and the differences between 3CT and 4CT spheroids. Finally, in the future, one might consider the involvement of additional cell types, such as other inflammatory cells, smooth muscle cells, or pericytes, to further define contraction and relaxation in in vitro models.

## Conclusion

This study presents a novel isogenic 3D spheroid model for investigating heart failure-associated stimuli. The integration of monocytes into 4CT spheroids significantly enhanced the model’s ability to replicate human heart tissue traits, as demonstrated by increased cardiomyocyte maturation and responsiveness to stimuli compared with 3CT spheroids. Although limitations such as small RNA-seq sample sizes and limited stimuli exist, this study lays a foundation for future research to explore additional mechanisms and stimuli, ultimately advancing heart failure research and personalised medicine.

## 6. Acknowledgements

We would like to thank Dr. K. Neef from the Regenerative Medicine Center Utrecht and T. Saric from the University of Cologne, Center for Physiology and Pathophysiology, Institute for Neurophysiology, Germany, for providing us with the iPSC lines. Figures were created with BioRender.com.

## Author Contributions

**E.T.S**.: Conceptualization, Methodology, Software, Formal analysis, Investigation, Resources, Data curation, Writing - Original Draft, Visualization. **E.L.K**.: Conceptualization, Methodology, Validation, Investigation, Resources, Writing - Original Draft, Visualization, Funding Acquisition. **E.F.v.V**.: Investigation, Writing - Original Draft, Visualization. **N.C**.: Investigation, Resources. **E.N**.: Investigation. **Q.C**.: Investigation. **A.v.M**.: Methodology, Validation, Writing - Review & Editing, Funding acquisition. **G.G.S**.: Writing - Review & Editing. **H.G**.: Writing - Review & Editing. **N.K**.: Writing - Review & Editing. **J.P.G.S**.: Conceptualization, Methodology, Resources, Writing - Review & Editing, Supervision, Funding acquisition. **S.d.J**.: Conceptualization, Methodology, Writing - Review & Editing, Supervision, Funding acquisition. **U.L**.: Conceptualization, Writing - Review & Editing, Supervision, Project administration, Funding acquisition.

## References

1. Timmis A, Vardas P, Townsend N, Torbica A, Katus H, De Smedt D, Gale CP, Maggioni AP, Petersen SE, Huculeci R, Kazakiewicz D, Benito Rubio V de, Ignatiuk B, Raisi-Estabragh Z, Pawlak A, Karagiannidis E, Treskes R, Gaita D, Beltrame JF, McConnachie A, Bardinet I, Graham I, Flather M, Elliott P, Mossialos EA, Weidinger F, Achenbach S, European Society of Cardiology, on behalf of the Atlas Writing Group. European Society of Cardiology: cardiovascular disease statistics 2021. Eur Heart J 2022;43:716–799.

2. DeBerge M, Shah SJ, Wilsbacher L, Thorp EB. Macrophages in Heart Failure with Reduced versus Preserved Ejection Fraction. Trends in Molecular Medicine 2019;25:328–340.

3. Lippi M, Stadiotti I, Pompilio G, Sommariva E. Human Cell Modeling for Cardiovascular Diseases. International Journal of Molecular Sciences 2020;21:6388.

4. Ham WB van, Kessler EL, Oerlemans MIFJ, Handoko ML, Sluijter JPG, Veen TAB van, Ruijter HM den, Jager SCA de. Clinical Phenotypes of Heart Failure With Preserved Ejection Fraction to Select Preclinical Animal Models. JACC: Basic to Translational Science 2022;7:844–857.

5. Drakhlis L, Biswanath S, Farr C-M, Lupanow V, Teske J, Ritzenhoff K, Franke A, Manstein F, Bolesani E, Kempf H, Liebscher S, Schenke-Layland K, Hegermann J, Nolte L, Meyer H, Roche J de la, Thiemann S, Wahl-Schott C, Martin U, Zweigerdt R. Human heart-forming organoids recapitulate early heart and foregut development. Nat Biotechnol 2021;39:737–746.

6. Ma C, Peng Y, Li H, Chen W. Organ-on-a-Chip: A New Paradigm for Drug Development. Trends in Pharmacological Sciences 2021;42:119–133.

7. Thomas D, Kim H, Lopez N, Wu JC. Fabrication of 3D Cardiac Microtissue Arrays using Human iPSC-Derived Cardiomyocytes, Cardiac Fibroblasts, and Endothelial Cells. J Vis Exp 2021.

8. Lewis-Israeli YR, Wasserman AH, Gabalski MA, Volmert BD, Ming Y, Ball KA, Yang W, Zou J, Ni G, Pajares N, Chatzistavrou X, Li W, Zhou C, Aguirre A. Self-assembling human heart organoids for the modeling of cardiac development and congenital heart disease. Nat Commun 2021;12:5142.

9. Zushin P-JH, Mukherjee S, Wu JC. FDA Modernization Act 2.0: transitioning beyond animal models with human cells, organoids, and AI/ML-based approaches. J Clin Invest 2023;133.

10. Directive 2010/63/EU of the European Parliament and of the Council of 22 September 2010 on the protection of animals used for scientific purposes (Text with EEA relevance)Text with EEA relevance

11. Alcaide P, Kallikourdis M, Emig R, Prabhu SD. Myocardial Inflammation in Heart Failure With Reduced and Preserved Ejection Fraction. Circulation Research 2024;134:1752–1766.

12. Kahn-Krell A, Pretorius D, Guragain B, Lou X, Wei Y, Zhang J, Qiao A, Nakada Y, Kamp TJ, Ye L, Zhang J. A three-dimensional culture system for generating cardiac spheroids composed of cardiomyocytes, endothelial cells, smooth-muscle cells, and cardiac fibroblasts derived from human induced-pluripotent stem cells. Front Bioeng Biotechnol 2022;10.

13. Jaén RI, Val-Blasco A, Prieto P, Gil-Fernández M, Smani T, López-Sendón JL, Delgado C, Boscá L, Fernández-Velasco M. Innate Immune Receptors, Key Actors in Cardiovascular Diseases. JACC: Basic to Translational Science 2020;5:735–749.

14. Lock RI, Graney PL, Tavakol DN, Nash TR, Kim Y, Sanchez E, Morsink M, Ning D, Chen C, Fleischer S, Baldassarri I, Vunjak-Novakovic G. Macrophages enhance contractile force in iPSC-derived human engineered cardiac tissue. Cell Reports 2024;43:114302.

15. Long C, Guo R, Han R, Li K, Wan Y, Xu J, Gong X, Zhao Y, Yao X, Liu J. Effects of macrophages on the proliferation and cardiac differentiation of human induced pluripotent stem cells. Cell Commun Signal 2022;20:108.

16. Matsa E, Burridge PW, Yu K-H, Ahrens JH, Termglinchan V, Wu H, Liu C, Shukla P, Sayed N, Churko JM, Shao N, Woo NA, Chao AS, Gold JD, Karakikes I, Snyder MP, Wu JC. Transcriptome Profiling of Patient-Specific Human iPSC-Cardiomyocytes Predicts Individual Drug Safety and Efficacy Responses In Vitro. Cell Stem Cell 2016;19:311–325.

17. Hamad S, Derichsweiler D, Papadopoulos S, Nguemo F, Šarić T, Sachinidis A, Brockmeier K, Hescheler J, Boukens BJ, Pfannkuche K. Generation of human induced pluripotent stem cell-derived cardiomyocytes in 2D monolayer and scalable 3D suspension bioreactor cultures with reduced batch-to-batch variations. Theranostics 2019;9:7222–7238.

18. Chirico N, Kessler EL, Maas RGC, Fang J, Qin J, Dokter I, Daniels M, Šarić T, Neef K, Buikema J-W, Lei Z, Doevendans PA, Sluijter JPG, Van Mil A. Small molecule-mediated rapid maturation of human induced pluripotent stem cell-derived cardiomyocytes. Stem Cell Res Ther 2022;13:531.

19. Lin Y, Linask KL, Mallon B, Johnson K, Klein M, Beers J, Xie W, Du Y, Liu C, Lai Y, Zou J, Haigney M, Yang H, Rao M, Chen G. Heparin Promotes Cardiac Differentiation of Human Pluripotent Stem Cells in Chemically Defined Albumin-Free Medium, Enabling Consistent Manufacture of Cardiomyocytes. Stem Cell Transl Med 2017;6:527–538.

20. Patsch C, Challet-Meylan L, Thoma EC, Urich E, Heckel T, O’Sullivan JF, Grainger SJ, Kapp FG, Sun L, Christensen K, Xia Y, Florido MHC, He W, Pan W, Prummer M, Warren CR, Jakob-Roetne R, Certa U, Jagasia R, Freskgård P-O, Adatto I, Kling D, Huang P, Zon LI, Chaikof EL, Gerszten RE, Graf M, Iacone R, Cowan CA. Generation of vascular endothelial and smooth muscle cells from human pluripotent stem cells. Nat Cell Biol 2015;17:994–1003.

21. Zhang J, Tao R, Campbell KF, Carvalho JL, Ruiz EC, Kim GC, Schmuck EG, Raval AN, Rocha AM da, Herron TJ, Jalife J, Thomson JA, Kamp TJ. Functional cardiac fibroblasts derived from human pluripotent stem cells via second heart field progenitors. Nat Commun 2019;10:2238.

22. Buchrieser J, James W, Moore MD. Human Induced Pluripotent Stem Cell-Derived Macrophages Share Ontogeny with MYB-Independent Tissue-Resident Macrophages. Stem Cell Rep 2017;8:334–345.

23. Yang X, Rodriguez M, Pabon L, Fischer KA, Reinecke H, Regnier M, Sniadecki NJ, Ruohola-Baker H, Murry CE. Tri-iodo-l-thyronine promotes the maturation of human cardiomyocytes-derived from induced pluripotent stem cells. Journal of Molecular and Cellular Cardiology 2014;72:296–304.

24. Friedman CE, Nguyen Q, Lukowski SW, Helfer A, Chiu HS, Miklas J, Levy S, Suo S, Han J-DJ, Osteil P, Peng G, Jing N, Baillie GJ, Senabouth A, Christ AN, Bruxner TJ, Murry CE, Wong ES, Ding J, Wang Y, Hudson J, Ruohola-Baker H, Bar-Joseph Z, Tam PPL, Powell JE, Palpant NJ. Single-Cell Transcriptomic Analysis of Cardiac Differentiation from Human PSCs Reveals HOPX-Dependent Cardiomyocyte Maturation. Cell Stem Cell 2018;23:586-598.e8.

25. Villani A-C, Satija R, Reynolds G, Sarkizova S, Shekhar K, Fletcher J, Griesbeck M, Butler A, Zheng S, Lazo S, Jardine L, Dixon D, Stephenson E, Nilsson E, Grundberg I, McDonald D, Filby A, Li W, De Jager PL, Rozenblatt-Rosen O, Lane AA, Haniffa M, Regev A, Hacohen N. Single-cell RNA-seq reveals new types of human blood dendritic cells, monocytes, and progenitors. Science 2017;356:eaah4573.

26. Hamidzada H, Pascual-Gil S, Wu Q, Kent GM, Massé S, Kantores C, Kuzmanov U, Gomez-Garcia MJ, Rafatian N, Gorman RA, Wauchop M, Chen W, Landau S, Subha T, Atkins MH, Zhao Y, Beroncal E, Fernandes I, Nanthakumar J, Vohra S, Wang EY, Valdman Sadikov T, Razani B, McGaha TL, Andreazza AC, Gramolini A, Backx PH, Nanthakumar K, Laflamme MA, Keller G, Radisic M, Epelman S. Primitive macrophages induce sarcomeric maturation and functional enhancement of developing human cardiac microtissues via efferocytic pathways. Nat Cardiovasc Res 2024;3:567–593.

27. O’Hern C, Caywood S, Aminova S, Kiselev A, Volmert B, Wang F, Sewavi M-L, Cao W, Dionise M, Muniyandi P, Popa M, Basrai H, Skoric M, Boulos G, Huang A, Nuñez-Regueiro I, Chalfoun N, Park S, Zhou C, Contag C, Aguirre A. Human heart assembloids with autologous tissue-resident macrophages recreate physiological immuno-cardiac interactions. bioRxiv 2024:2024.11.13.623051.

28. Suku M, Murphy JF, Corbezzolo S, Biggs M, Forte G, Turnbull IC, Costa KD, Forrester L, Monaghan MG. Synergistic generation of cardiac resident-like macrophages and cardiomyocyte maturation in tissue engineered platforms

29. Peters MC, Maas RGC, Adrichem I van, Doevendans PAM, Mercola M, Šarić T, Buikema JW, Mil A van, Chamuleau SAJ, Sluijter JPG, Hnatiuk AP, Neef K. Metabolic Maturation Increases Susceptibility to Hypoxia-induced Damage in Human iPSC-derived Cardiomyocytes. Stem Cells Transl Med 2022;11:1040–1051.

30. Mishra S, Kass DA. Cellular and molecular pathobiology of heart failure with preserved ejection fraction. Nat Rev Cardiol 2021:1–24.

31. Haim IR, Gruber A, Kazma N, Bashai C, Lichtig Kinsbruner H, Caspi O. Modeling Heart Failure With Preserved Ejection Fraction Using Human Induced Pluripotent Stem Cell-Derived Cardiac Organoids. Circ Heart Fail 2025:e011690.

32. Somers T, Siddiqi S, Maas RGC, Sluijter JPG, Buikema JW, Broek PHH van den, Meuwissen TJ, Morshuis WJ, Russel FGM, Schirris TJJ. Statins affect human iPSC-derived cardiomyocytes by interfering with mitochondrial function and intracellular acidification. Basic Res Cardiol 2024;119:309–327.

33. Maas RGC, Beekink T, Chirico N, Snijders Blok CJB, Dokter I, Sampaio-Pinto V, Mil A van, Doevendans PA, Buikema JW, Sluijter JPG, Stillitano F. Generation, High-Throughput Screening, and Biobanking of Human-Induced Pluripotent Stem Cell-Derived Cardiac Spheroids. J Vis Exp 2023.

34. Azarisman SM, Teo KS, Worthley MI, Worthley SG. Cardiac magnetic resonance assessment of diastolic dysfunction in acute coronary syndrome. J Int Med Res 2017;45:1680–1692.

35. Inoue T, Iseki K, Iseki C, Ohya Y, Kinjo K, Takishita S. Effect of heart rate on the risk of developing metabolic syndrome. Hypertens Res 2009;32:801–806.

36. Zile MR, Baicu CF, Gaasch WH. Diastolic Heart Failure — Abnormalities in Active Relaxation and Passive Stiffness of the Left Ventricle. New England Journal of Medicine 2004;350:1953–1959.

37. Henein MY, Lindqvist P. Assessment of Left Ventricular Diastolic Function by Doppler Echocardiography. 2015.

38. Nagueh SF. Left Ventricular Diastolic Function: Understanding Pathophysiology, Diagnosis, and Prognosis With Echocardiography. JACC: Cardiovascular Imaging 2020;13:228–244.

39. Packer M, Lam CSP, Lund LH, Maurer MS, Borlaug BA. Characterization of the inflammatory-metabolic phenotype of heart failure with a preserved ejection fraction: a hypothesis to explain influence of sex on the evolution and potential treatment of the disease. European Journal of Heart Failure 2020;22:1551–1567.

40. Hedman ÅK, Hage C, Sharma A, Brosnan MJ, Buckbinder L, Gan L-M, Shah SJ, Linde CM, Donal E, Daubert J-C, Mälarstig A, Ziemek D, Lund L. Identification of novel pheno-groups in heart failure with preserved ejection fraction using machine learning. Heart 2020;106:342–349.

41. Potere N, Bonaventura A, Abbate A. Novel Therapeutics and Upcoming Clinical Trials Targeting Inflammation in Cardiovascular Diseases. Arteriosclerosis, Thrombosis, and Vascular Biology 2024;44:2371–2395.

42. Kessler EL, Oerlemans MIFJ, Hoogen P van den, Yap C, Sluijter JPG, Jager SCA de. Immunomodulation in Heart Failure with Preserved Ejection Fraction: Current State and Future Perspectives. J Cardiovasc Transl 2021;14:1–12.

43. Peh ZH, Dihoum A, Hutton D, Arthur JSC, Rena G, Khan F, Lang CC, Mordi IR. Inflammation as a therapeutic target in heart failure with preserved ejection fraction. Front Cardiovasc Med 2023;10.

44. Mooney L, Jackson CE, Adamson C, McConnachie A, Welsh P, Myles RC, McMurray JJV, Jhund PS, Petrie MC, Lang NN. Adverse Outcomes Associated With Interleukin-6 in Patients Recently Hospitalized for Heart Failure With Preserved Ejection Fraction. Circulation: Heart Failure 2023;16:e010051.

45. Castillo EC, Vázquez-Garza E, Yee-Trejo D, García-Rivas G, Torre-Amione G. What Is the Role of the Inflammation in the Pathogenesis of Heart Failure? Curr Cardiol Rep 2020;22:139.

46. Mil A van, Balk GM, Neef K, Buikema JW, Asselbergs FW, Wu SM, Doevendans PA, Sluijter JPG. Modelling inherited cardiac disease using human induced pluripotent stem cell-derived cardiomyocytes: progress, pitfalls, and potential. Cardiovasc Res 2018;114:1828–1842.

47. Kawaguchi N, Nakanishi T. Stem Cell Studies in Cardiovascular Biology and Medicine: A Possible Key Role of Macrophages. Biology 2022;11:122.

48. Incalza MA, D’Oria R, Natalicchio A, Perrini S, Laviola L, Giorgino F. Oxidative stress and reactive oxygen species in endothelial dysfunction associated with cardiovascular and metabolic diseases. Vasc Pharmacol 2018;100:1–19.

49. McHugh K, DeVore AD, Wu J, Matsouaka RA, Fonarow GC, Heidenreich PA, Yancy CW, Green JB, Altman N, Hernandez AF. Heart Failure With Preserved Ejection Fraction and Diabetes JACC State-of-the-Art Review. J Am Coll Cardiol 2019;73:602–611.

50. Anker SD, Butler J, Filippatos G, Ferreira JP, Bocchi E, Böhm M, Rocca H-PB-L, Choi D-J, Chopra V, Chuquiure-Valenzuela E, Giannetti N, Gomez-Mesa JE, Janssens S, Januzzi JL, Gonzalez-Juanatey JR, Merkely B, Nicholls SJ, Perrone SV, Piña IL, Ponikowski P, Senni M, Sim D, Spinar J, Squire I, Taddei S, Tsutsui H, Verma S, Vinereanu D, Zhang J, Carson P, Lam CSP, Marx N, Zeller C, Sattar N, Jamal W, Schnaidt S, Schnee JM, Brueckmann M, Pocock SJ, Zannad F, Packer M, Investigators E-PT. Empagliflozin in Heart Failure with a Preserved Ejection Fraction. New Engl J Med 2021.

