## Supplement for "Isogenic monocytes improve the responsiveness of hiPSC cardiac spheroids to cardiac stressors"

### Supplementary Figures

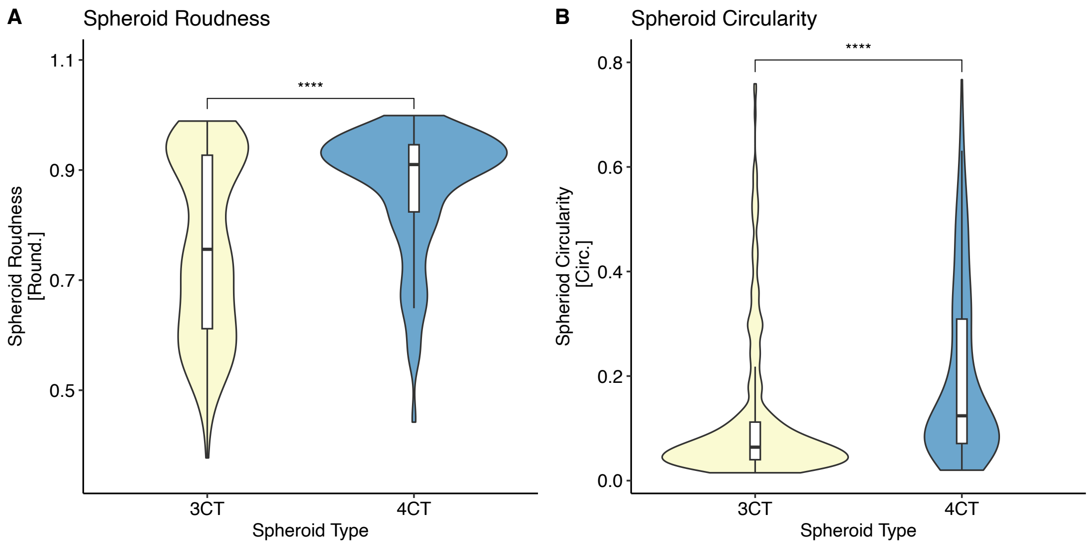


**Supplementary Figure S1. Analysis of roundness and circularity of spheroids. A)** 4CT spheroids are significantly rounder [calculated as: $\frac{4\pi(\mathrm{Area})}{Major\ Axis^2}$] than 3CT spheroids (p<0.0001). **B)** 4CT spheroids are also significantly more circular [calculated as: $\frac{4\pi(\mathrm{Area})}{Perimeter^2}$ ] than 3CT spheroids (p<0.0001). Circularity is especially sensitive to blebbing, whereas roundness compares how close a shape is to a perfect circle based on perimeter and area. Statistical analysis: Wilcoxon rank test. n = 18 per group.

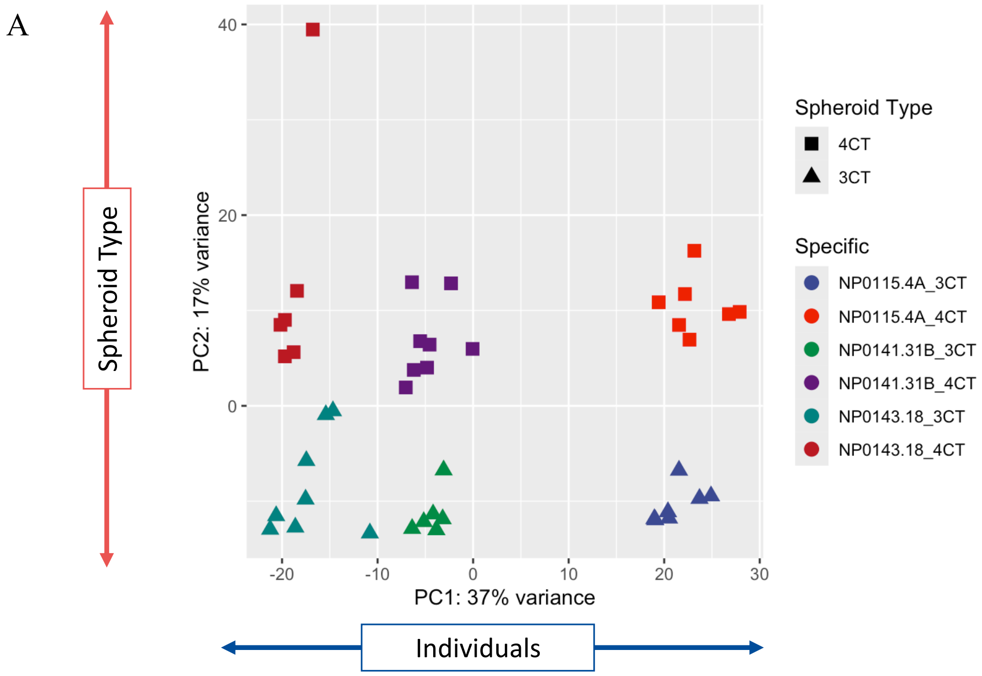

B

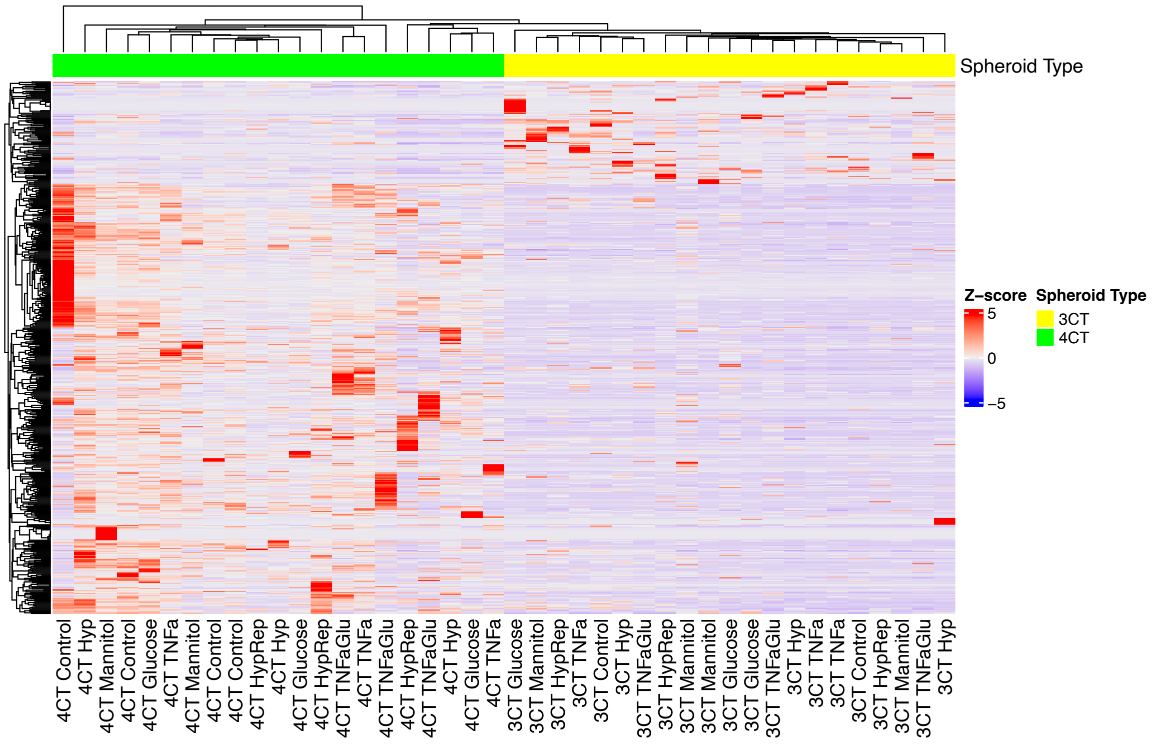
Spheroid Type
Z-score
5
0
−5
Spheroid Type
3CT
4CT
4CT Control
4CT Hyp
4CT Mannitol
4CT Control
4CT Glucose
4CT TNFa
4CT Mannitol
4CT Control
4CT Control
4CT HypRep
4CT Hyp
4CT Glucose
4CT HypRep
4CT TNFaGlu
4CT TNFa
4CT TNFaGlu
4CT HypRep
4CT TNFaGlu
4CT Hyp
4CT Glucose
4CT TNFa
3CT Glucose
3CT Mannitol
3CT HypRep
3CT TNFa
3CT Control
3CT Hyp
3CT TNFaGlu
3CT HypRep
3CT Mannitol
3CT Mannitol
3CT Glucose
3CT Glucose
3CT TNFaGlu
3CT Hyp
3CT TNFa
3CT TNFa
3CT Control
3CT HypRep
3CT Mannitol
3CT TNFaGlu
3CT Hyp

C

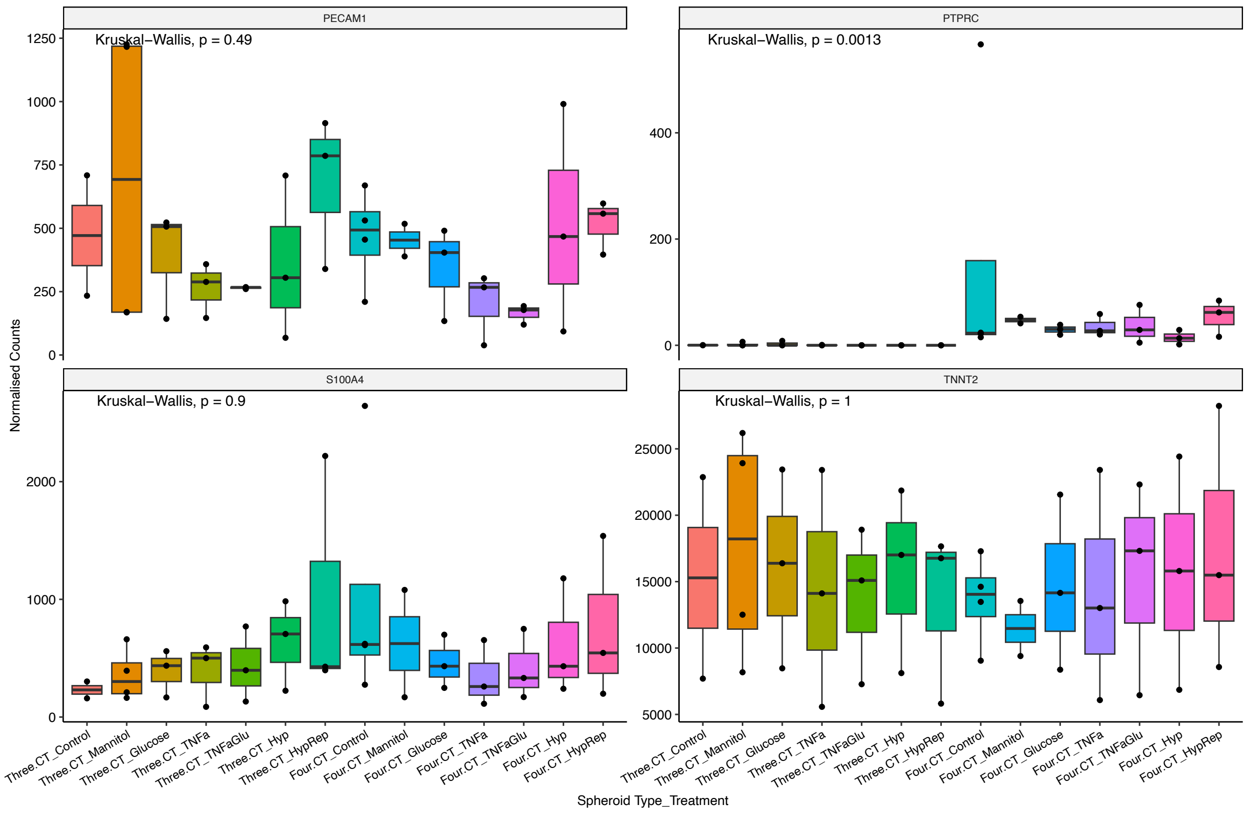
PECAM1
Kruskal–Wallis, p = 0.49
PTPRC
Kruskal–Wallis, p = 0.0013
S100A4
Kruskal–Wallis, p = 0.9
TNNT2
Kruskal–Wallis, p = 1
Normalised Counts
Spheroid Type_Treatment
Three.CT_Control
Three.CT_Mannitol
Three.CT_Glucose
Three.CT_TNFa
Three.CT_TNFaGlu
Three.CT_Hyp
Three.CT_HypRep
Four.CT_Control
Four.CT_Mannitol
Four.CT_Glucose
Four.CT_TNFa
Four.CT_TNFaGlu
Four.CT_Hyp
Four.CT_HypRep

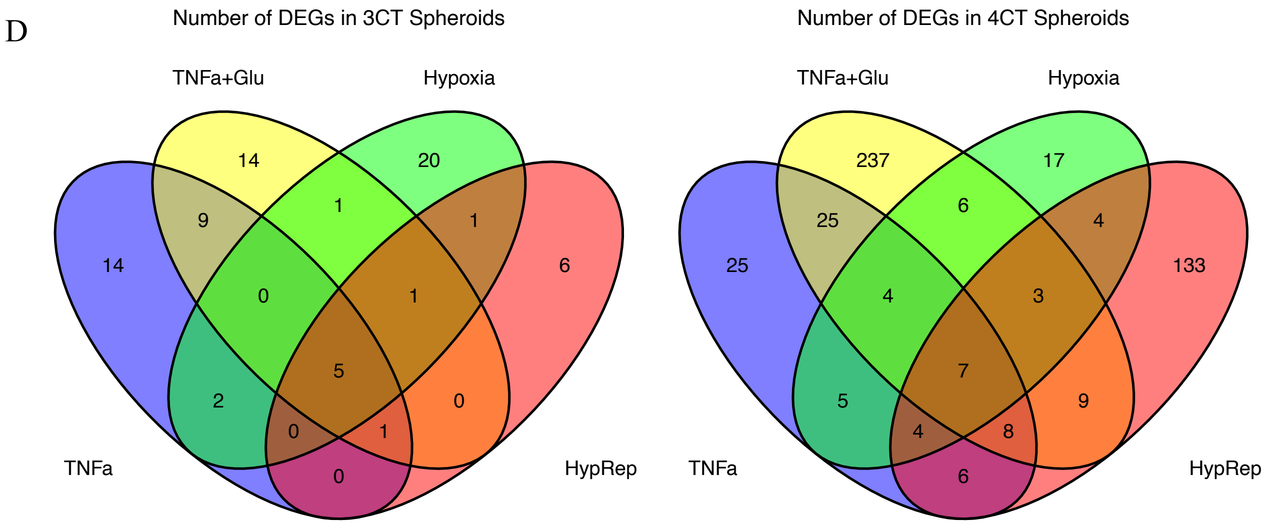


**Supplementary Figure S2. RNA-sequencing analysis. A)** Plot of principal component analysis of RNA-sequencing data. **B)** Heat map of RNA-sequencing data. **C)** Normalized counts of cell type markers in several types and treatments of cardiac spheroids. There is no significant difference between groups in cell type marker expression. PECAM1 for endothelial cells, PPTRC (CD45) for monocytes/macrophages, S100A4 for fibroblasts and TNNT2 for cardiomyocytes. **D)** The number of differentially expressed genes in 3CT (left) and 4CT (right) spheroids depending on treatment (cutoff: adjusted p-value < 0.05 & log2 fold change > 0.6).

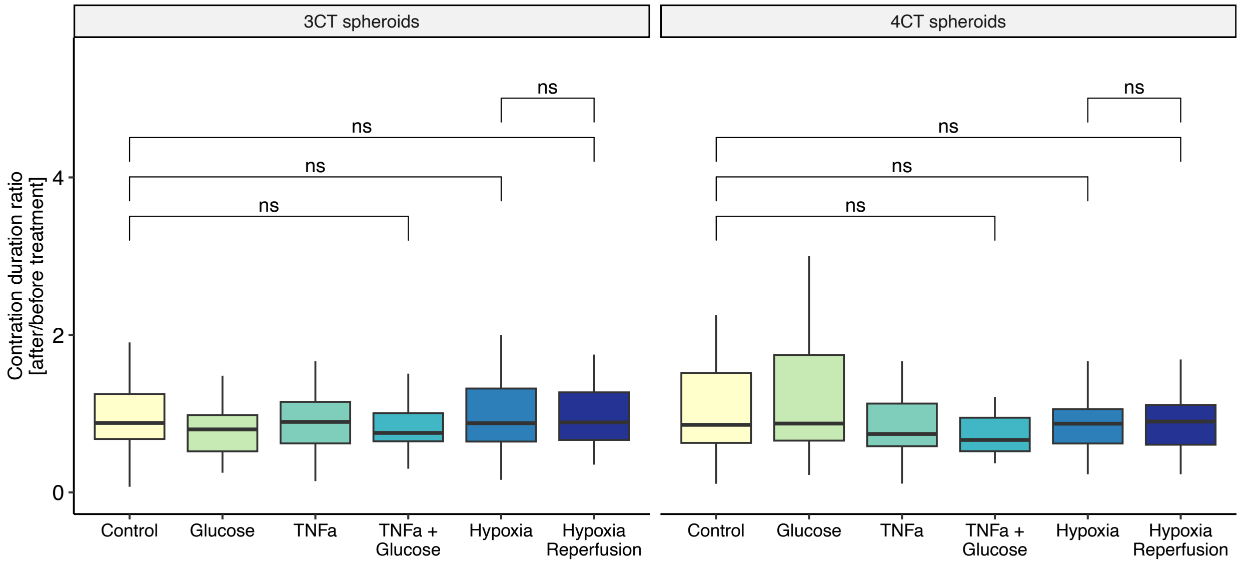

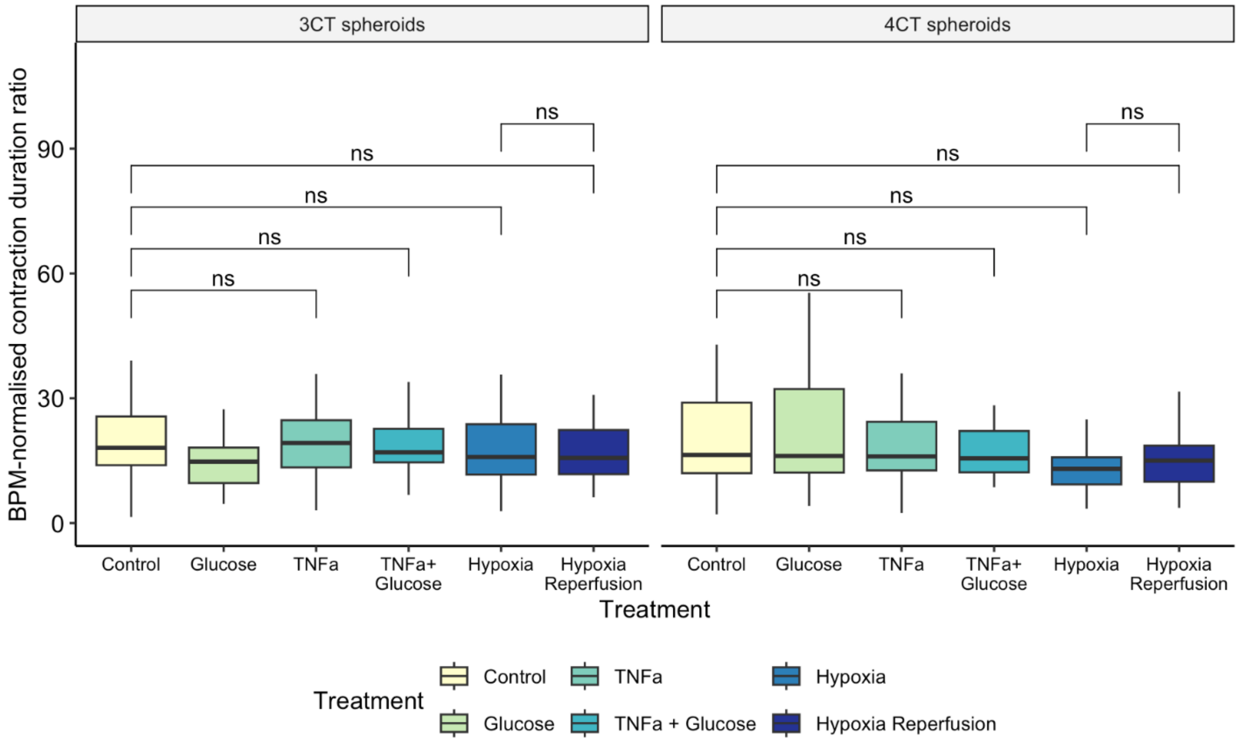


**Supplementary Figure S3. Contraction duration ratio and normalized contraction duration ratio.**

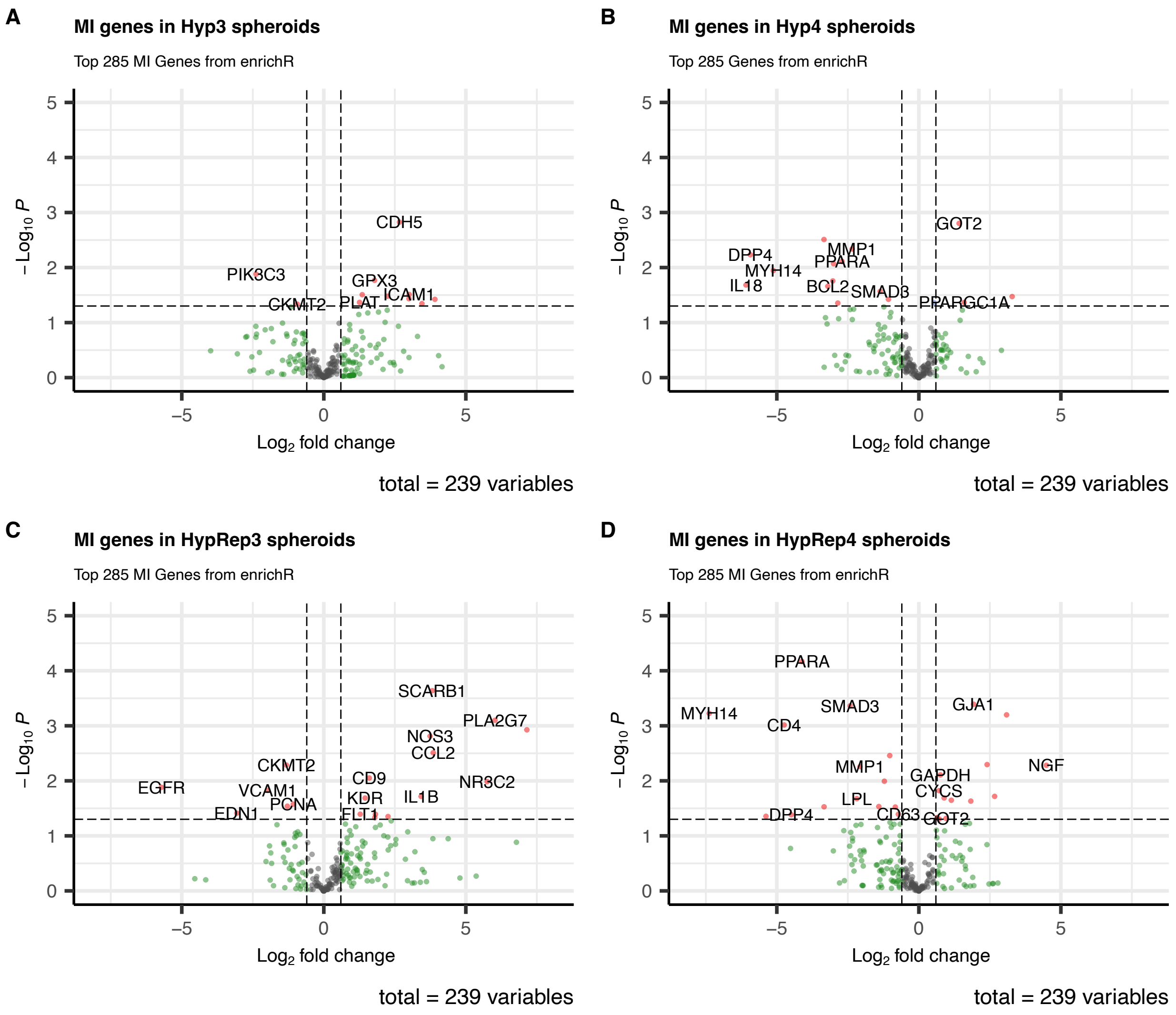


**Supplementary Figure S4. Differentially expressed MI-associated genes in 3CT (left) and 4CT (right) spheroids in hypoxia (0.5% for 6h) treated vs control spheroids and reoxygenated hypoxia.**

**Supplemental Tables**: List of significantly differentially expressed HFpEF-associated genes.

Differentially expressed HFpEF genes in 3CT under TNFa & glucose treatment

| | *baseMean* | *log2FoldChange* | *lfcSE* | *stat* | *pvalue* | *p.adj* |
|---|---|---|---|---|---|---|
| *HLA-C* | 734.323662 | 3.02117175 | 0.78831614 | 3.83243674 | 0.00012688 | 0.00748593 |
| *TAPBP* | 323.617802 | 2.12795946 | 0.66913459 | 3.1801666 | 0.0014719 | 0.02470273 |
| *TCF3* | 131.665995 | 1.90981033 | 0.67427575 | 2.83238769 | 0.00462018 | 0.03894151 |
| *AP1M1* | 181.494235 | 1.76244101 | 0.56082932 | 3.14256219 | 0.00167476 | 0.02470273 |
| *DDX54* | 45.1027418 | 2.36289823 | 0.71804981 | 3.29071631 | 0.00099933 | 0.02470273 |
| *TSC2* | 327.61319 | -1.9012429 | 0.62913638 | -3.0219885 | 0.0025112 | 0.02872062 |
| *DGKQ* | 12.5888367 | 5.17445916 | 1.73875362 | 2.97595881 | 0.00292074 | 0.02872062 |

Differentially expressed HFpEF genes in 4CT under TNFa & glucose treatment

| | *baseMean* | *log2FoldChange* | *lfcSE* | *stat* | *pvalue* | *p.adj* |
|---|---|---|---|---|---|---|
| *RBM38* | 910.367754 | -2.5266882 | 0.69563485 | -3.6322048 | 0.00028101 | 0.00319649 |
| *ANKRD11* | 201.797165 | -3.1370752 | 0.66168758 | -4.7410218 | 2.13E-06 | 0.00019351 |
| *KANK2* | 282.392438 | -1.8038737 | 0.5249993 | -3.4359545 | 0.00059047 | 0.00510771 |
| *EMILIN1* | 138.842396 | 3.59304326 | 0.87727285 | 4.09569639 | 4.21E-05 | 0.0019151 |
| *CAPN1* | 104.747499 | -1.5709766 | 0.56905868 | -2.7606584 | 0.0057685 | 0.02101305 |
| *ESAM* | 464.362891 | -1.9059191 | 0.50056668 | -3.8075229 | 0.00014037 | 0.00255466 |
| *GRN* | 1769.18763 | -1.7188745 | 0.46335289 | -3.7096446 | 0.00020755 | 0.00283867 |
| *NME3* | 100.935911 | 1.72334025 | 0.53361846 | 3.22953639 | 0.00123991 | 0.00723171 |
| *MYH14* | 51.7403428 | -7.5130295 | 2.10771846 | -3.5645318 | 0.00036451 | 0.00368556 |
| *TK1* | 158.908171 | -1.6693574 | 0.61947148 | -2.6948091 | 0.0070429 | 0.02373717 |
| *SEMA6C* | 34.2493345 | -5.8136369 | 1.47686882 | -3.9364612 | 8.27E-05 | 0.00192425 |
| *CCDC85B* | 80.5166742 | -2.0235712 | 0.77943275 | -2.5962101 | 0.00942584 | 0.02599247 |
| *NOTCH1* | 120.56076 | -2.4786586 | 0.78812142 | -3.1450212 | 0.00166075 | 0.00888989 |
| *MGLL* | 42.8137834 | -4.1756376 | 1.22888309 | -3.3979128 | 0.00067902 | 0.00514924 |
| *ZMIZ2* | 99.8651514 | 1.722619 | 0.65996028 | 2.61018588 | 0.0090493 | 0.02589321 |
| *ACE* | 23.9291644 | -4.0133496 | 1.2864316 | -3.1197536 | 0.00181002 | 0.00915068 |
| *KIF26A* | 28.8158996 | -2.4224551 | 0.92017069 | -2.6326149 | 0.00847304 | 0.02589321 |
| *ROBO4* | 23.0240351 | -3.2176074 | 1.09838097 | -2.9294093 | 0.00339607 | 0.01475232 |
| *NPDC1* | 175.466955 | -1.7057114 | 0.61791868 | -2.760414 | 0.00577282 | 0.02101305 |
| *SH2D3C* | 317.269839 | -4.7956061 | 1.21993609 | -3.9310306 | 8.46E-05 | 0.00192425 |
| *RABL6* | 186.573782 | -1.7800102 | 0.58225191 | -3.0571135 | 0.0022348 | 0.0107035 |

| | | | | | | |
|---|---|---|---|---|---|---|
| *LAMA5* | 205.982305 | -1.7890338 | 0.61087329 | -2.9286495 | 0.00340438 | 0.01475232 |
| *CAPN15* | 169.729328 | -2.4551781 | 0.76192542 | -3.2223339 | 0.00127151 | 0.00723171 |
| *ZNF768* | 41.6817104 | -3.2113173 | 1.14463773 | -2.8055316 | 0.00502337 | 0.01987506 |
| *RAMP1* | 65.7954049 | -2.8049357 | 0.75875297 | -3.6967707 | 0.00021836 | 0.00283867 |
| *PLCD3* | 73.7146498 | -2.0980382 | 0.89677663 | -2.3395326 | 0.01930789 | 0.04765471 |
| *NMRK2* | 55.5018297 | -3.333188 | 0.97352159 | -3.423846 | 0.00061742 | 0.00510771 |
| *PPP1R13L* | 53.6916868 | 2.49934955 | 0.95299028 | 2.62263909 | 0.00872516 | 0.02589321 |
| *MACROD1* | 32.4556145 | -2.1608384 | 0.92414011 | -2.3382151 | 0.01937609 | 0.04765471 |
| *F2RL3* | 33.2645957 | -3.9483979 | 1.1924707 | -3.3111069 | 0.00092928 | 0.00650494 |
| *NOTCH4* | 88.0225143 | -3.2647156 | 1.00628999 | -3.2443089 | 0.00117736 | 0.00723171 |
| *ZNF414* | 38.2442675 | 2.02928086 | 0.77807617 | 2.60807482 | 0.00910531 | 0.02589321 |
| *STAB1* | 39.826427 | -2.6274998 | 1.0223555 | -2.5700452 | 0.01016853 | 0.02721576 |
| *RILP* | 98.3715684 | 1.72729373 | 0.6483201 | 2.66426068 | 0.00771577 | 0.02507627 |
| *CROCC* | 9.52428108 | -4.1196038 | 1.70808171 | -2.4118306 | 0.01587265 | 0.0412689 |
| *RABEP2* | 13.1550265 | -3.6944688 | 1.34758101 | -2.741556 | 0.00611489 | 0.02140213 |
| *IFI27* | 39.5295278 | -4.5500079 | 1.57875462 | -2.8820235 | 0.0039513 | 0.01634402 |

Differentially expressed HFpEF genes in 3CT under hypoxia treatment

| | *baseMean* | *log2FoldChange* | *lfcSE* | *stat* | *pvalue* | *p.adj* |
|---|---|---|---|---|---|---|
| *NFIC* | 936.825912 | 1.85780268 | 0.72958973 | 2.54636626 | 0.01088509 | 0.0467271 |
| *WDR13* | 119.510353 | 2.40862722 | 0.79577683 | 3.0267622 | 0.00247188 | 0.02449462 |
| *CTDSP1* | 256.761692 | 1.50444737 | 0.58009025 | 2.59347123 | 0.00950125 | 0.0467271 |
| *SYNPO* | 214.627418 | -2.1723342 | 0.72958442 | -2.9774953 | 0.00290614 | 0.02449462 |
| *AP1M1* | 181.494235 | 1.58470903 | 0.56245554 | 2.8174832 | 0.00484016 | 0.0347196 |
| *MLST8* | 61.3142284 | -3.059953 | 0.72584745 | -4.2156971 | 2.49E-05 | 0.00073457 |
| *CDH5* | 181.426123 | 2.6630245 | 0.83945493 | 3.17232577 | 0.00151223 | 0.02230543 |
| *SIVA1* | 113.251685 | -1.7898979 | 0.70178164 | -2.5505055 | 0.01075668 | 0.0467271 |
| *C1QTNF1* | 140.94776 | -3.8617967 | 1.39485735 | -2.7685962 | 0.00562984 | 0.0347196 |
| *MEGF8* | 43.4968286 | 3.40209639 | 1.13404842 | 2.99995689 | 0.00270018 | 0.02449462 |
| *ARFRP1* | 58.469082 | 6.30408123 | 1.20175076 | 5.24574766 | 1.56E-07 | 9.18E-06 |
| *NUDT22* | 103.771757 | 1.80262006 | 0.70971531 | 2.53991992 | 0.01108779 | 0.0467271 |
| *DGKQ* | 12.5888367 | 5.61424484 | 1.7475626 | 3.21261444 | 0.00131533 | 0.02230543 |
| *CDK18* | 13.3145444 | 3.07890463 | 1.11791913 | 2.75413895 | 0.00588468 | 0.0347196 |

Differentially expressed HFpEF genes in 4CT under hypoxia treatment

| | *baseMean* | *log2FoldChange* | *lfcSE* | *stat* | *pvalue* | *p.adj* |
|---|---|---|---|---|---|---|
| *RBM38* | 910.367754 | -2.7829182 | 0.6931181 | -4.0150707 | 5.94E-05 | 0.00273369 |
| *HLA-B* | 771.557899 | -1.8040424 | 0.72273709 | -2.4961254 | 0.01255582 | 0.02764462 |
| *ANKRD11* | 201.797165 | -1.6909808 | 0.62047468 | -2.7253019 | 0.00642427 | 0.0254317 |
| *ARSA* | 31.2350636 | 2.55051702 | 0.86738271 | 2.94047483 | 0.0032771 | 0.02171082 |
| *ARHGEF10L* | 81.732887 | 1.75872704 | 0.63631725 | 2.7639154 | 0.00571123 | 0.0254317 |
| *MYH14* | 51.7403428 | -5.1221423 | 2.02349118 | -2.5313391 | 0.01136279 | 0.02750992 |
| *CLDN5* | 159.781655 | -1.8363282 | 0.75728554 | -2.4248822 | 0.01531336 | 0.02938912 |
| *ZNHIT2* | 20.6078474 | -2.0836919 | 0.84343057 | -2.470496 | 0.01349258 | 0.02821176 |
| *MGLL* | 42.8137834 | -2.9170376 | 1.15163486 | -2.5329535 | 0.0113106 | 0.02750992 |
| *ACE* | 23.9291644 | -3.0019852 | 1.14009234 | -2.633107 | 0.00846077 | 0.02594636 |
| *ROBO4* | 23.0240351 | -2.5168093 | 0.94915185 | -2.6516403 | 0.00801018 | 0.02594636 |
| *CAPN15* | 169.729328 | -2.0130375 | 0.74154274 | -2.7146614 | 0.00663436 | 0.0254317 |
| *AHDC1* | 43.5991569 | 1.80309632 | 0.82572724 | 2.18364639 | 0.02898824 | 0.04938737 |
| *C19orf24* | 59.454911 | -1.7270723 | 0.69240595 | -2.494306 | 0.01262037 | 0.02764462 |
| *ABCD1* | 98.6564378 | -1.9342226 | 0.65835618 | -2.9379578 | 0.00330382 | 0.02171082 |
| *AP5Z1* | 46.3485827 | -2.0512043 | 0.78674372 | -2.6072077 | 0.0091284 | 0.02624414 |
| *INTS1* | 78.2929579 | -1.797935 | 0.67942179 | -2.6462723 | 0.00813843 | 0.02594636 |
| *NMRK2* | 55.5018297 | -1.879116 | 0.83261019 | -2.2568977 | 0.02401447 | 0.04418662 |
| *PPP1R13L* | 53.6916868 | 3.30082486 | 0.93907883 | 3.51496036 | 0.00043982 | 0.00924149 |
| *F2RL3* | 33.2645957 | -3.1423879 | 1.04847093 | -2.997115 | 0.00272548 | 0.02171082 |
| *NOTCH4* | 88.0225143 | -2.9131242 | 0.95582289 | -3.0477657 | 0.0023055 | 0.02171082 |
| *SIGIRR* | 63.2875538 | -1.8772957 | 0.77433224 | -2.4244059 | 0.01533346 | 0.02938912 |
| *KMT2B* | 50.1949876 | -2.6161296 | 0.91942632 | -2.8453934 | 0.00443566 | 0.0254317 |
| *MROH1* | 34.5807309 | -3.5084346 | 1.0227498 | -3.4303938 | 0.00060271 | 0.00924149 |
| *ZNF710* | 39.4329676 | 2.16651089 | 0.77860147 | 2.78256717 | 0.00539307 | 0.0254317 |
| *NARFL* | 18.9182887 | -1.8704801 | 0.85633909 | -2.184275 | 0.02894205 | 0.04938737 |
| *MLPH* | 8.07390106 | -5.9797892 | 2.36132222 | -2.5323902 | 0.01132879 | 0.02750992 |

Differentially expressed HFpEF genes in 3CT under hypoxia reoxygenation treatment

| | *baseMean* | *log2FoldChange* | *lfcSE* | *stat* | *pvalue* | *p.adj* |
|---|---|---|---|---|---|---|
| *HLA-C* | 734.323662 | 2.96188676 | 0.78826073 | 3.75749627 | 0.00017162 | 0.00892434 |

Differentially expressed HFpEF genes in 4CT under hypoxia reoxygenation treatment

| | *baseMean* | *log2FoldChange* | *lfcSE* | *stat* | *pvalue* | *p.adj* |
|---|---|---|---|---|---|---|
| *PCNX3* | 39.7003124 | -2.292413297 | 0.9726316 | -2.3569184 | 0.0184273 | 0.04054005 |
| *RBM38* | 910.3677538 | -2.99571323 | 0.69640539 | -4.3016801 | 1.70E-05 | 0.00037292 |
| *SCOC* | 310.1420046 | 1.98547313 | 0.53364169 | 3.72061097 | 0.00019874 | 0.00218616 |
| *ANKRD11* | 201.7971654 | -1.583435242 | 0.63332796 | -2.5001821 | 0.01241295 | 0.03034276 |
| *RGS3* | 468.6888603 | 2.324942068 | 0.58673692 | 3.96249493 | 7.42E-05 | 0.00097905 |
| *KLHL21* | 165.7293823 | 1.859762181 | 0.69052536 | 2.6932569 | 0.00707577 | 0.02030439 |
| *MYH14* | 51.7403428 | -7.047490424 | 2.08933149 | -3.3730839 | 0.00074331 | 0.00490587 |
| *NOTCH3* | 151.7429844 | 2.233484514 | 0.8133998 | 2.74586311 | 0.00603519 | 0.01810558 |
| *ARHGEF17* | 80.55354116 | 1.868460543 | 0.67010616 | 2.78830526 | 0.00529846 | 0.01770638 |
| *MYO1C* | 652.9441986 | 1.94490316 | 0.56187402 | 3.46145774 | 0.00053726 | 0.00443238 |
| *ATN1* | 219.7362036 | 1.574056414 | 0.57115282 | 2.75592862 | 0.00585258 | 0.01810558 |
| *MGLL* | 42.81378341 | -3.831025754 | 1.21001234 | -3.1661047 | 0.00154495 | 0.0078436 |
| *ROBO4* | 23.0240351 | -2.499957724 | 0.97370512 | -2.567469 | 0.01024439 | 0.02817208 |
| *SLC27A1* | 39.21367329 | 3.596744324 | 1.1290223 | 3.18571592 | 0.00144396 | 0.0078436 |
| *SH2D3C* | 317.269839 | -3.361428695 | 1.2073119 | -2.7842256 | 0.00536557 | 0.01770638 |
| *RABL6* | 186.573782 | -2.037106475 | 0.58213695 | -3.4993595 | 0.00046638 | 0.00439727 |
| *CAPN15* | 169.7293282 | -2.294234479 | 0.76508798 | -2.9986545 | 0.00271175 | 0.00994307 |
| *CIC* | 60.40399474 | 2.999290766 | 0.88596488 | 3.38533822 | 0.00071091 | 0.00490587 |
| *OBSL1* | 604.9305309 | 3.00187783 | 0.64124999 | 4.68129107 | 2.85E-06 | 9.41E-05 |
| *SLC4A3* | 102.5535785 | 1.564359385 | 0.61886882 | 2.52777216 | 0.01147888 | 0.0291387 |
| *TOLLIP* | 139.8544305 | 2.412646681 | 0.606017 | 3.98115344 | 6.86E-05 | 0.00097905 |
| *ABHD8* | 87.10346547 | 4.807382105 | 0.86890159 | 5.53271185 | 3.15E-08 | 2.08E-06 |
| *PLCD3* | 73.71464984 | -2.268878317 | 0.89694475 | -2.5295631 | 0.01142047 | 0.0291387 |
| *SPG7* | 104.7540013 | 1.928857984 | 0.62894732 | 3.06680373 | 0.00216361 | 0.00892489 |
| *NOTCH4* | 88.0225143 | -3.37507403 | 1.01090183 | -3.3386763 | 0.00084179 | 0.00505072 |
| *PRR12* | 14.39893912 | -3.256561737 | 1.04659923 | -3.1115652 | 0.00186098 | 0.00818833 |
| *KMT2B* | 50.19498763 | -2.217479438 | 0.95096106 | -2.3318299 | 0.01970964 | 0.04196246 |
| *MROH1* | 34.5807309 | -2.354868189 | 1.03511079 | -2.2749914 | 0.02290644 | 0.04724454 |
| *KRBA1* | 38.43838701 | -2.277498472 | 0.9215495 | -2.4713794 | 0.01345929 | 0.03172548 |
| *MYH7B* | 115.1340306 | 1.916769352 | 0.78360758 | 2.44608322 | 0.01444177 | 0.03286747 |
| *POLD1* | 22.71294561 | 3.029424731 | 1.00551277 | 3.01281577 | 0.00258836 | 0.00994307 |
| *ACACB* | 36.77673382 | 2.840015781 | 0.90966313 | 3.12205222 | 0.00179595 | 0.00818833 |
